# Structure of Photosystem I-FCP from giant kelp uncovers drivers of antenna evolution across the red lineage

**DOI:** 10.1101/2025.10.07.680316

**Authors:** J. D Weissman, P. Maturana, H. M. O. Oung, R. Riddle, G. Wyatt, V. G. T. Dubinin, P. Zerbe, M. Maldonado

## Abstract

Brown algae and other red-algae-derived organisms such as diatoms are major contributors to global CO_2_ fixation *via* photosynthesis. To understand the photosynthetic function of brown algae, we obtained the structure of giant kelp *Macrocystis pyrifera* photosystem I (PSI) with a fucoxanthin-chlorophyll-protein (FCP) antenna and compared it to known structures from the red-algal lineage. We identified differences in *M. pyrifera*’s antenna composition, architecture and chlorophyll networks, as well as a pronounced variation in transmembrane hydrophobic thickness across the PSI-FCP supercomplex, with implications for photochemical function. Our work lays the foundation to understand kelp’s high photosynthetic productivity, reveals new drivers of antenna conservation and diversification, and sheds light on evolutionary relationships between organisms of the red lineage.

## Main Text

The photosynthetic conversion of inorganic carbon into organic material, i.e., primary production, sustains most life on this planet (*1*). Roughly half of the global primary production originates from photosynthesis in the oceans, carried out by a range of prokaryotic and eukaryotic organisms (*2*). A diverse group of eukaryotic phototrophs derived from red algae (“red lineage”) are major contributors to primary production, marine ecosystem maintenance and coastal economic activity around the world (*2–5*). Understanding the precise molecular mechanisms of photosynthesis and how it has diversified across the red lineage will illuminate this critical metabolic process and enable the development of new ocean-based environmental strategies. Here, we focus on the structural features of the photosynthetic complex photosystem I (PSI) of brown algae to shed light on its specialization and diversification, as well as on the evolutionary relationships between the different clades of the red lineage.

Oxygenic photosynthesis uses solar energy to convert carbon dioxide (CO_2_) and water into organic molecules and molecular oxygen. The photochemical reactions of photosynthesis are enabled by multi-protein photosystems I and II (PSI, PSII) in the thylakoid membrane, which harness light to power the transfer of electrons from water to NADP^+^ and pump protons into the thylakoid lumen. PSI catalyzes the transfer of electrons from electron donors such as plastocyanin (Pc) and cytochrome *c*_6_ to the electron acceptor ferredoxin (Fd). The synthesis of NAPDH and ATP resulting from the light reactions of photosynthesis drives the fixation of carbon from CO_2_ into organic molecules (*1*).

Photosystems absorb light by the excitation of chromophores such as chlorophylls (chl). To increase the efficiency of light absorption and to obtain protection against photo-oxidative damage, in many organisms photosystems associate with additional light-harvesting proteins that form a multi-protein “antenna” (*6*). The antenna funnels the absorbed light into the photosystem reaction center by transferring the energy across the chromophores in a process known as excitation energy transfer (EET) (*1*). Whereas photosystems are structurally and functionally conserved, antenna systems and the chromophores they contain are highly diverse, tuned to optimal absorption of wavelengths in different environments (*6*). The most abundant antenna system in eukaryotic phototrophs, including the red lineage, is that composed of light-harvesting complex (LHC) membrane proteins. The LHC family has diversified into subfamilies that share a three-transmembrane-helix topology but can differ in the number and types of their bound chromophores (*6*, *7*). The LHC diversification is salient in the red lineage, where the Lhcr subfamily present in red algae expanded and evolved into at least six different subfamilies in subsequent clades (*6*, *7*). Additionally, LHCs in the red lineage have been historically referred to by their bound chromophores, e.g., fucoxanthin-chlorophyll *a*/*c* protein (FCP), regardless of their LHC subfamily (*6–8*).

Red algae, as well as plants and green algae, originated from the engulfment of a photosynthetic cyanobacterium. Through gene transfers and other processes, this primary endosymbiont evolved into a stable organelle, i.e., the chloroplast, of the red alga (*1*, *9*). The red lineage includes the diverse phyla Cryptophyta, Haptophyta, Ochrophyta (e.g., photosynthetic stramenopiles such as diatoms and brown algae) and Myzozoa (e.g., dinoflagellates) (**fig. S1A-C**). In these phyla, higher-order endosymbiosis of a red alga or another red-lineage organism led to the formation of increasingly complex chloroplasts upon engulfment and stabilization through gene transfer. Although it is clear that the red lineage did *not* originate from a single secondary endosymbiosis of a red alga (*10–12*), the evolutionary trajectory of the set remains elusive, with phylogenomic support for various models of serial or parallel endosymbioses (*11–15*) (**fig. S1A-D**). Understanding the evolutionary relationships between the red-lineage phyla and the evolution of their photosystems and their antenna is key to understand this critical group of primary producers.

The comparative study of protein structure and subunit composition provides a complementary approach to disambiguate evolutionary relationships between organisms, especially with multi-protein complexes involved in core metabolism, such as PSI (*16*, *17*). Currently, high-resolution structures of PSI with its antenna assembly are available for various red algae, cryptophytes, haptophytes and dinoflagellates (*18–27*). Among ochrophytes, structures are only available for diatoms, organisms of the Diatomista clade (*28–31*) (**fig. S1E**). Although diatoms are major marine phototrophs and well-studied model systems, their features should not be taken as representative of the entire ochrophyte phylum (*32*) (**fig. S1E**). For instance, several features identified in diatom PSI are lacking non-Diatomista ochrophytes at the sequence level (*30*). The structural study of an ochrophyte of the Chrysista clade, e.g., a brown alga, would resolve questions about the generalizability of features across the phylum and the red lineage.

To tackle these questions, we determined the structure and composition of a PSI-FCP antenna supercomplex assembly of brown alga *Macrocystis pyrifera* (giant kelp). Kelp forests are critical components of coastal ecosystems and economies, exhibiting primary productivity in line with terrestrial crops and up to three times higher than marine phytoplankton (*3–5*). Thus, kelp forests could provide significant additions to ocean-based climate solutions, e.g., *via* carbon sequestration (*33*, *34*). Understanding the molecular details of brown algal photosynthesis is fundamental to develop applications that leverage kelp’s photosynthetic abilities.

Through detailed comparisons of the *M. pyrifera* PSI-FCP structure with the rest of the red lineage, we identified differences between Chrysista and Diatomista EET pathways, with implications for photosynthetic function. We identified membrane-thickness variation across PSI-FCPs of red-lineage organisms and propose that hydrophobic mismatch regulates the function of the supercomplex. We posit that contingent protein:protein interactions, gene loss and neutral evolution are key drivers of antenna conservation and diversification across the red lineage and discuss how structural comparisons can help elucidate evolutionary relationships between the phyla of the red lineage.

### *M. pyrifera* PSI-FCP supercomplex

After optimizing protocols to decrease the viscosity of the kelp sample, we isolated a chloroplast-enriched fraction from fresh *M. pyrifera* blades. We used digitonin extraction and sucrose gradient ultracentrifugation to purify PSI-FCP and proceeded to structural determination using cryogenic electron microscopy (cryoEM) (**figs. S2-S6**, **table S1-S3**). We observed a PSI-FCP supercomplex with 11 PSI subunits (PsaA/B/C/D/E/F/I/J/L/M/R) and 18 FCP subunits arranged in two belts (first belt: FCP1-11; second belt: FCP13/15/16/17/19/A/B) (**Fig. 1, figs. S3-S6**). We followed the FCP numbering of diatom *Chaetoceros gracilis*, the largest observed ochrophyte antenna, to minimize re-naming (see **fig. S7** for conversion) (*28*). Note that FCP number refers to the position in the antenna, *not* to the gene name. The structure also models 3 [4Fe-4S] clusters, 2 phylloquinones, 218 chl *a*, 27 chl *c*, 23 β-carotenes, 26 violaxanthins, 19 fucoxanthins and 18 lipids (**fig. S2**, **tables S3-S4**). We detected and quantified the pigments chlorophyll *a*, chlorophyll *c*, fucoxanthin, violaxanthin, zeaxanthin and β-carotene using high performance liquid chromatography (**fig. S2**).

**Fig. 1.**
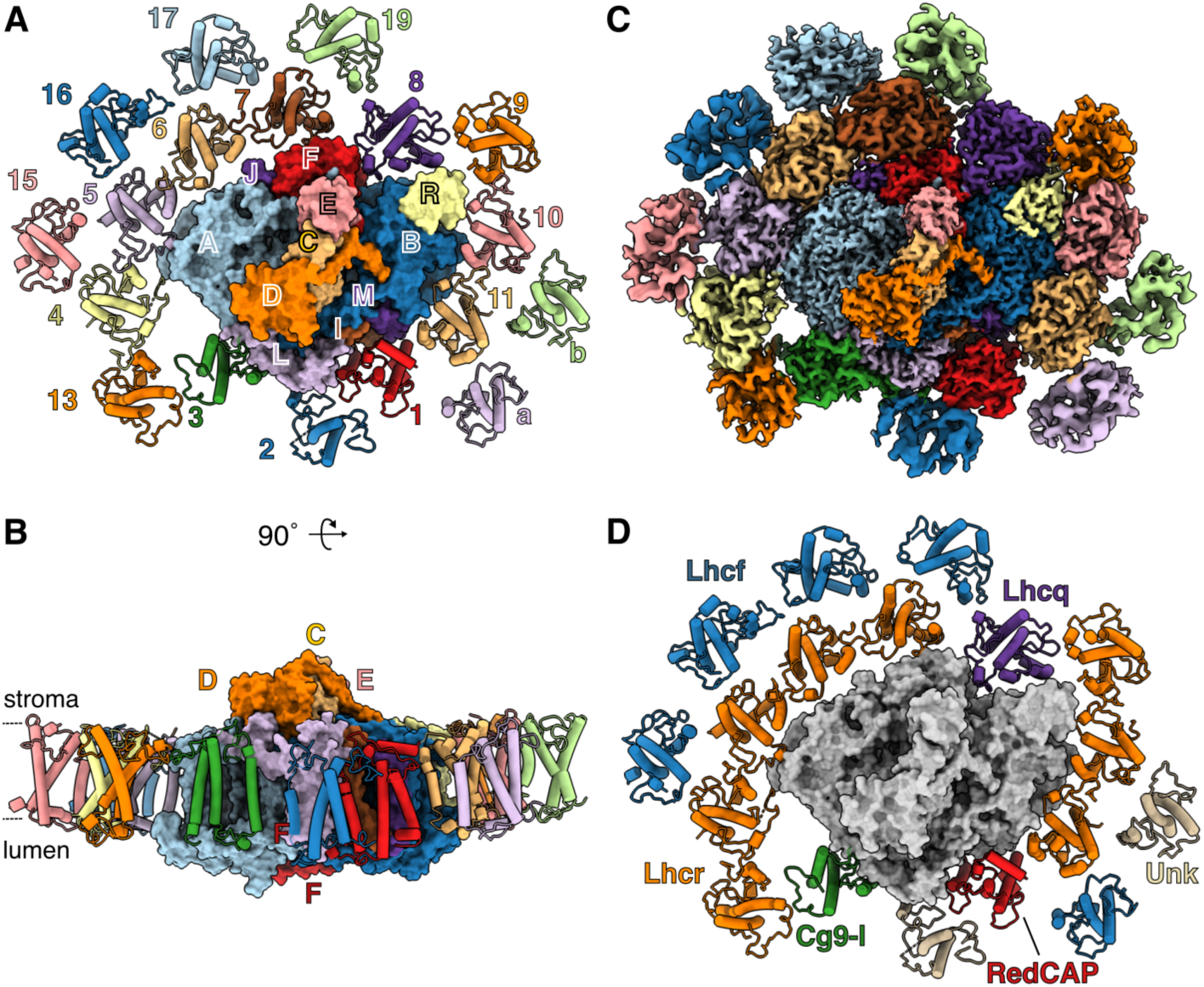
Structure of *Macrocystis pyrifera*’s PSI-FCP supercomplex. (**A-B**) Atomic model of *M. pyrifera* PSI-FCP supercomplex viewed from the stroma (A) or the membrane (B). PSI in surface, FCP proteins in cartoon, colored by subunit. The approximate locations of the stroma and lumen are shown with dashed lines. For clarity, “Psa” and “FCP” suffixes are omitted from subunit labels (e.g., PsaA labeled A, FCP labeled 1). To prevent confusion with PsaA/B, FCPA/B are labeled in small print. (**C**) Stromal view of cryoEM composite map of *M. pyrifera* PSI-FCP supercomplex, colored by subunit as in (A-B). (**D**) Stromal view of *M. pyrifera* PSI-FCP supercomplex labeled by FCP subfamily. PSI shown in grey; Lhcr in orange; RedCAP in red; CgLhcr9-like (Cg9-l) in green; Lhcq in purple; Lhcf in blue; unknown family (Unk) in beige.

Overall, protein, chromophore and electron transfer co-factor components of PSI exhibited high structural conservation with previous red-lineage structures (*18–31*). Inspection of the plastocyanin, cytochrome *c*_6_ and ferredoxin binding regions of PSI revealed almost complete conservation (*35*) (**Supplementary Text, fig. S8**). We did not find evidence for stromal subunit PsaS, which was recently reported in diatoms (*28*, *30*), in either the cryoEM density or the *M. pyrifera* genome (*36*). This supports the notion that PsaS is a Diatomista innovation (*30*). The structure also confirmed the lack of PsaK in brown algal PSI (present in red algae, cryptophytes and haptophytes), as well as the lack of PsaO (present in red algae and cryptophytes) (*18–31*).

### *M. pyrifera* FCP phylogeny

To increase our understanding of FCP diversification in the red lineage, we identified *M. pyrifera*’s FCP complement in the genome and EST databases and performed phylogenetic analyses using previously identified LHC protein sequences from diatoms and brown algae (*37*, *38*, *36*) (**fig. S9, Supplementary Data File 1**). The resulting tree resolved the 57 *M. pyrifera* FCPs into seven subfamilies (Lhcr, Lhcf, Lhcq, Lhcx, Lhcz, CgLhcr9-like and RedCAP), with Lhcz as a subclade or sister clade to Lhcr as seen in diatoms (*38*) (**fig. S9**). This further establishes that this phylogenetic classification is robust and common across the red lineage (*38*, *39*). Whereas orthology was evident in subfamilies Lhcr, CgLhcr9-like and Lhcq, we observed substantial species-specific expansion of subfamilies Lhcf and Lhcx (**fig. S9**). Despite the size of the Lhcx subfamily (eight genes in *M. pyrifera*; six in diatom *Thalassiosira pseudonana*, **fig. S9**), only one member has been reported in a diatom PSII structure (*40*), and none in PSI (*18–31*). Lhcx proteins play photoprotective roles in the energy-dependent component of non-photochemical quenching *via* PSII in diatoms (*41–43*). Given that *M. pyrifera* does not use this type of photoprotection (*44*, *45*), the presence of eight Lhcx genes suggests they may have an alternative function and/or be retained by neutral selection.

### Structure of *M. pyrifera* FCP antenna

All FCP subunits showed the expected topology (transmembrane helices B, C, A) and Arg-Glu salt bridges between helices A and B seen in all photosynthetic organisms so far (*6*, *46*) (**figs. S10A-F)**. Chromophore-binding motifs identified in diatoms for each FCP subfamily are also conserved in *M. pyrifera* (*38*) (**fig. S10G-J**). As seen in other red-lineage organisms, FCP interactions were mediated mostly on the stromal side, by FCPs’ N-terminal loop and helix C regions (FCP:PSI, **fig. S10K,L**), or N-terminal loop and C-A loop (FCP:FCP, **fig. S10M,N**) (*18*–*31*).

To elucidate functionally relevant differences between FCP subfamilies and between organisms, we performed detailed comparisons of the antennae across the red lineage (**figs. S10-S11**). For clarity, all antenna proteins are referred to as “FCP” using our numbering, even if they bind other chromophores in certain organisms (e.g., dinoflagellates’ peridin-chlorophyll-*a*-proteins). Based on our phylogenetic analysis, *M. pyrifera*’s first belt was composed of 11 FCP subunits (FCP1-11) of the Lhcr, Lhcq, CgLhcr9-like subfamilies and RedCAP family (**figs. S9-S11**). The second belt was composed of FCP13/15/16/17/19/A/B, which belong to the Lhcr and Lhcf subfamilies (**figs. S9-S11**). Of the 18 FCP subunits observed in our structure, FCPA/B were new positions not previously seen in other organisms (*18–31*) (**Fig. 1, fig. S11**). Using a biochemical preparation with a different digitonin ratio, we observed *M. pyrifera* PSI-FCP supercomplexes with larger antennae suggestive of additional new positions in the vicinity of FCPA/B (**fig. S12**). The size and shape of these larger *M. pyrifera* antenna were different from those of the currently largest antennae of diatoms or haptophytes (*25*, *28*) (**fig. S12**). However, the resolution of these maps was insufficient for atomic modeling.

### M. pyrifera’s first belt: Lhcr majority affects membrane thickness

The majority of *M. pyrifera*’s first belt FCPs belong to subfamily Lhcr (FCP4/5/6/7/9/10/11, **Fig. 1D**), as seen across all organisms of the red lineage (*18–31*) (**fig. S11**). The majority is absolute in red algae and cryptophytes (*18–21*, *38*, *39*) (**table S5**). We posit that the first-belt Lhcr majority is driven by conserved interactions with PSI subunits, as well as by structural features of the subfamily (see **Discussion**).

Contacts between the first belt and PSI proteins, as well as between FCPs across the antenna, are mediated by various types of hydrophobic interactions on the stromal side (**Fig. 2A-C, fig. S10K-N**). Lhcr proteins in the first belt establish hydrophobic interactions between aromatic residues on the N-terminus and the C-A loop of the binding partners (**Fig. 2A-B)**. These aromatic-mediated interactions are specific to the Lhcr family and conserved across organisms (*18–31*). Interactions between *M. pyrifera*’s Lhcr:Lhcr pairs in the first belt showed a more positive energy of dissociation relative to Lhcr binding to other subfamilies, indicating more stable interactions (average ΔG dissociation 6.75 kcal/mol vs −8.81 kcal/mol, respectively). Moreover, in *M. pyrifera* and others, Lhcr proteins harbor a subfamily-specific chromophore-binding sequence motif: SxS/AL/IP in the N-terminal loop for chl 415 binding (*38*) (**fig. S10G**). Lhcrs’ chl 415 is ∼6 Å away from chl 407 of the adjacent FCP, poising these chlorophyll molecules for efficient energy transfer into the core (**Fig. 2C-E**). Indeed, fast transfer (<10 ps) between chl 415 and chl 407 of Lhcr subunits in the first belt has been measured in dinoflagellates (*27*). Thus, replacing Lhcr proteins in the first belt with other subfamilies leads to changes to the EET network.

**Fig. 2.**
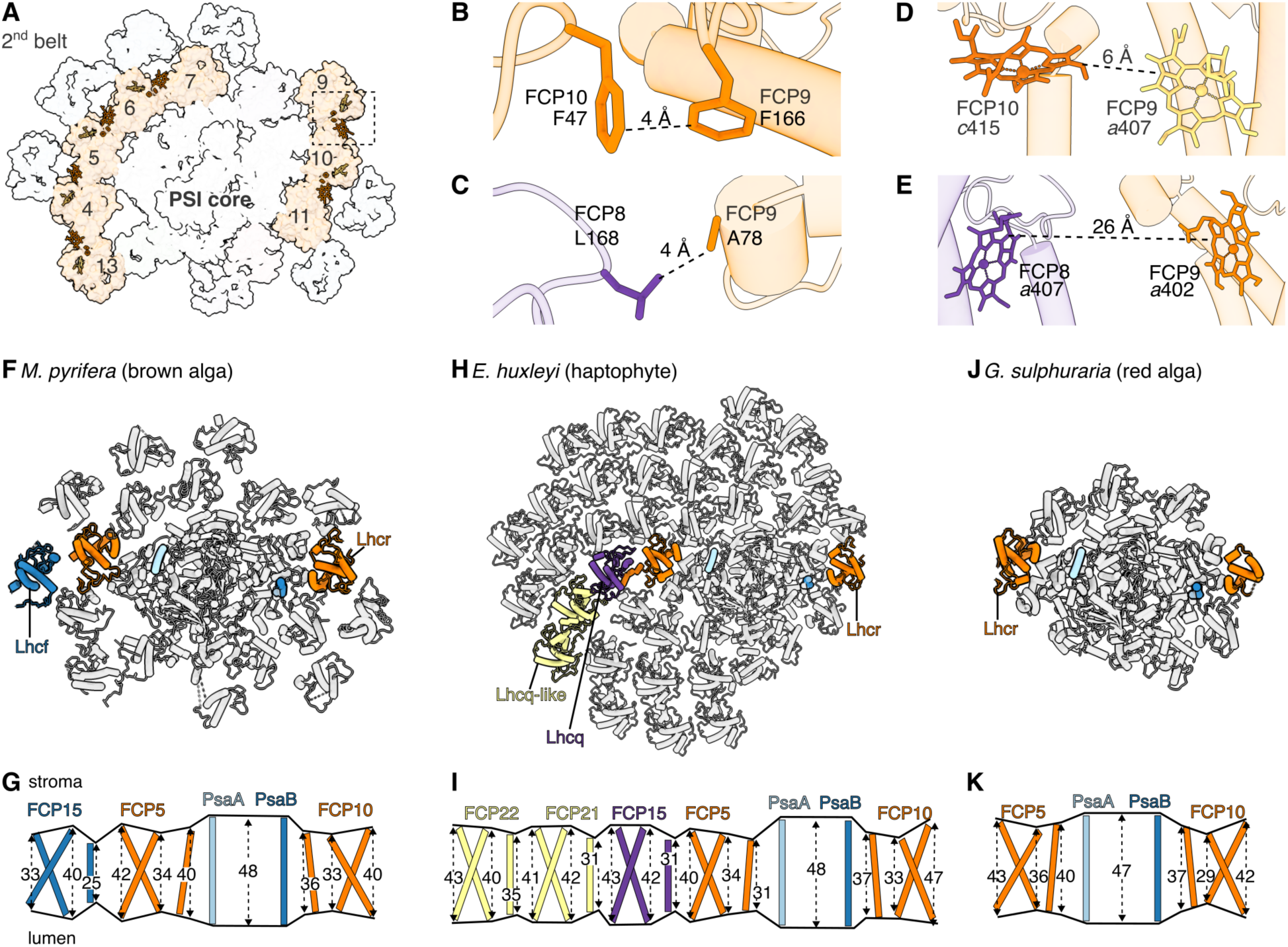
FCP structural features affect interactions and membrane thickness of PSI-FCP belts. (**A**) Stromal view of *M. pyrifera* PSI-FCP highlighting Lhcr subunits and features discussed in text. Lhcr subunits shown in orange surface with labels with FCP number; core and second belt in grey surface. Porphyrin ring of key chlorophyll molecules and benzene ring of key phenylalanine residues shown in atom representation. Lhcr-specific features shown in dark orange; Lhcr chlorophyll found in other families shown in yellowish-orange. Dashed box indicates area detailed in (B-C). (**B-E**) Details of hydrophobic interactions (B-C) and chlorophyll-mediated energy transfer (D-E) between an Lhcr pair (B,D) or an Lhcr-non-Lhcr pair (C,E). Components are colored by their respective FCP subfamilies (Lhcr in orange, Lhcq in purple). Edge-to-edge distances between interaction partners indicated. **(F-K)** Membrane thickness profiles of PSI-FCP from *M. pyrifera* (F,G) (PDB: 9YGV, this work), *Emiliania huxleyi* (H, I) (PDB: 9JJ8) (*25*), *Galdieria sulphuraria* (J, K) (PDB: 9KC5) (*18*). (F-J) Stromal views, subunits in cartoon representation. For best fit, (H) is scaled to 80% relative to (F,J). Subunits with helices thicknesses highlighted in (G-K) shown colored by FCP subfamily (Lhcr in orange, Lhcq in purple, Lhcf in blue, Lhcq-like in yellow); helices from PsaA in light blue, from PsaB in dark blue. (G-K) Schematic of membrane thickness of subunits colored in (F-J). Membrane thickness was estimated from the height of the helices in the respective models, measured in angstrom and indicated by dashed arrows. The changes in membrane thickness are marked by a black trace of the helix spans.

Additionally, we noticed differences in the FCP helix-C across subfamilies. In the *M. pyrifera* structure, Lhcr FCPs possess the longest helices C (largest number of amino acids) (**fig. S10**) as well as the largest height in the membrane, with an average of 31.5 Å (**Fig. 2G**). Given than Lhcr helices C are usually bent and somewhat tilted, their height is smaller than would otherwise be calculated for a straight helix. The Lhcr height contrasts with subfamily averages of 26.1 Å, 27.2 Å, 24.5 Å and 23.8 Å for first-belt members of the RedCAP, Cg9-l, Lhcq and Lhcf subfamilies, respectively (**Supplementary Data File 2**). The largest observed difference was of 16 Å and 10 residues between Lhcr-FCP5 (40 Å, 28 residues) and Lhcf-FCP17 (24 Å, 18 residues, **Fig. 2G, Supplementary Data File 2)**.

We also observed differences in helices C across the first and second belt, with averages of 30.1 Å (Lhcr majority) and 24.8 Å (proposed Lhcf majority, see below, **Fig. 2F-G**). Furthermore, helices A and B are longer than helix C across all FCPs (averaging 39.5 Å and 33.2 Å respectively), and PSI subunits PsaA/PsaB house the longest helices in the supercomplex, spanning 47.6 Å on average (**Fig. 2G**). Given that most FCPs are oriented such that helix C is placed between helices A/B of two FCPs of the previous belt, the positioning of B/A-C-B/A-C helices creates a radially alternating pattern of helix heights (**Fig. 2G**). Together, these observations suggest both a rippling and overall thinning of the membrane from PSI towards the periphery of the antenna (**Fig. 2G**).

This rippling and thinning pattern is present in other red lineage PSI-FCP supercomplexes, albeit to different extents (**Fig. 2H-K**). In haptophyte *Emiliania huxleyi*, the largest red-lineage PSI-FCP described to date, PsaA/B remain the tallest region at 47.8 Å. Inferred membrane thickness then shows slight fluctuations across FCP subfamilies Lhcr, Lhcq, and Lhcq-like, with helix-C averages of 33.3 Å, 30.5 Å, and 30.7 Å respectively in subsequent antenna belts. Helices A/B average 37.8 and 40.1 Å across FCP subfamilies (**Fig. 2H-I**). The rippling effect is the least pronounced in ancestral-like red algae *Galdieria sulphuraria*, with its unanimous Lhcr FCP composition and lack of second belt (**Fig. 2J-K**). Nevertheless, the thickest points of *G. sulphuraria* PSI are analogous to *M. pyrifera*, spanning 47 Å. The red alga shows a thinning in the first belt, with helix-C of Lhcr FCPs averaging 38 Å (**Fig. 2J-K**).

These local differences in membrane thickness have functional implications for the photochemical properties of chromophores at different antenna locations (*47–50*). Additionally, membrane-thickness differences may lead to in hydrophobic mismatch, which we hypothesize could be exploited to regulate PSI-FCP function (see **Discussion**) (*51–57*). Spectroscopic and biophysical experiments, as well as in situ imaging of *M. pyrifera* chloroplast membranes using cryo-electron tomography will shed light on these topics.

### MpFCP3: rotations and family switches enabled by loss of PsaO

In contrast to the Lhcr majority, in *M. pyrifera* the FCP3 position is occupied by a CgLhcr9-like (Cg9-l) subfamily protein (**Fig. 1D**, **fig. S11)**. Thus, *M. pyrifera*’s FCP3 lacks the Lhcr protein-binding and ligand motif otherwise seen in the first belt (**fig. S10G**). Moreover, *M. pyrifera*’s FCP3 shows a ∼135° rotation along its vertical axis relative to other first-belt FCPs and to cryptophyte FCP3, which is an Lhcr protein (**Fig. 3A-B**). This rotation exposes the FCP3’s C-A loop and the “backside” of helix C towards the core, placing the N-terminus towards the second belt. The new interfaces allow FCP3 to form hydrophobic and electrostatic interactions with core-subunit PsaL (**Fig. 3A-D**). The rotation also changes the location of *M. pyrifera* FCP3’s chromophores, decreasing the chl distances to PsaL from ∼16 Å seen in cryptophytes to ∼6 Å in *M. pyrifera* (*22*, *23*) (**Fig. 3E-F**).

**Fig. 3.**
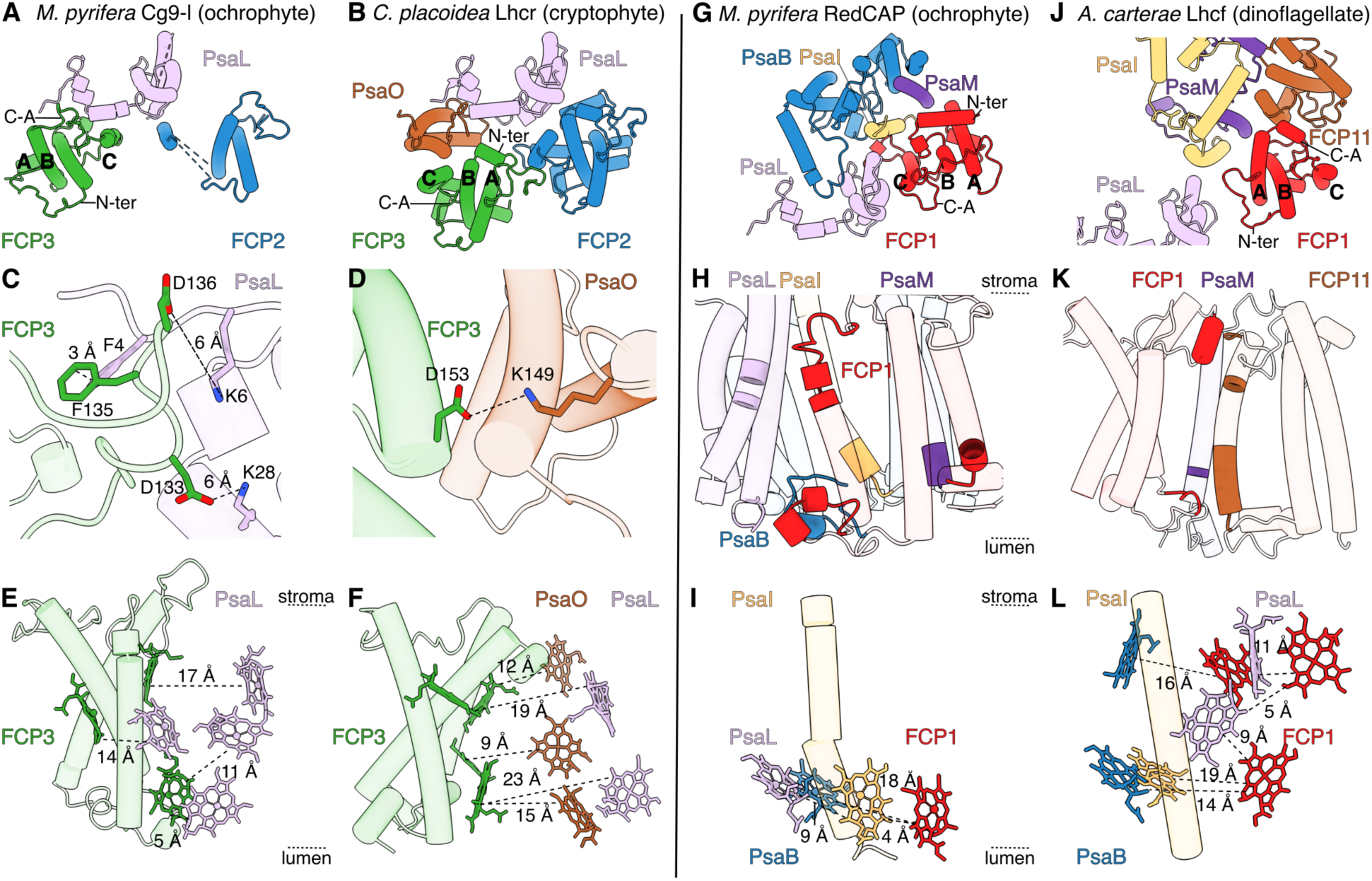
FCP subfamily switches for FCP3 and FCP1. (**A-F**) FCP3 switch from Lhcr to CgLhr9-like (Cg9-l) subfamily. (**G-L**) FCP1 switch from RedCAP to Lhcf subfamily. (A, C, E, G-I) Details for *M. pyrifera* (brown alga, ochrophyte, PDB: 9YGV, this work). (B, D, F) Details for *Croomonas placoidea* (cyroptophyte, PDB: 7Y7B) (*22*). (J-L) Details for *Amphidinium cartera* (dinoflagellate, PDB: 8JW0) (*26*). (A, B, G, H) Relevant subunits discussed in the text, viewed from the stroma, shown in cartoon, colored by subunit. Helices A, B and C, C-A loop (C-A) and N-terminal loop (N-ter) labeled. (C, D, H, K) Details of key protein:protein interactions viewed from the stroma (C, D) or from the membrane (H, K). (E, F, I, L) Interactions of key chlorophyll molecules between the partners in (C, D, H, K) viewed from the membrane. Edge-to-edge distances between the porphyrin rings shown. Approximate placement of stroma and lumen shown with dashed lines.

An analogous FCP3 rotation and subfamily switch is seen in diatoms, haptophytes and dinoflagellates (**fig. S11**). In contrast, FCP3 remains an Lhcr protein in cryptophytes (*22*, *23*). FCP3 is lacking in the available red algal structures (*20*, *21*, *19*, *18*). If this is due to loss during purification, it is reasonable to assume that red algal FCP3 is also Lhcr, since that is the only subfamily present (*38*, *39*). Given this, as well as the fact that cryptophytes do not possess the Cg9-l subfamily (*38*), the ancestral state can be assumed to be an Lhcr FCP3.

We propose that the changes in FCP3 family and orientation can be explained by changes in its ancestral interaction partner PsaO, which is encoded in the nucleus of red algae and cryptophytes and lost in the rest of the lineage (*24*, *26–30*, *58*, *59*). The presence of PsaO correlates with an Lhcr FCP3 in standard orientation; the loss of PsaO (ochrophytes, haptophytes, dinoflagellates) perfectly correlates with a Cg9-l FCP3 in rotated orientation. We hypothesize that the loss of PsaO is the result of an endosymbiotic engulfment of a cryptophyte, followed by a failure to transfer PsaO from the nucleomorph to the nucleus of the new host before the nucleomorph was degraded. The loss of PsaO produced a physical gap in PSI and broke contingent interactions between PSI and the Lhcr FCP3, allowing for the replacement by a new family in a new orientation, with re-configured EET connections (**Fig. 3E-F**).

### MpFCP1: extensive RedCAP-PSI protein interactions with weak EET

*M. pyrifera*’s FCP1 belongs to the RedCAP family of the LHC superfamily (*60*). RedCAP is the most structurally distinct protein in the red-lineage antennae, with an extended helix F in the N-terminal loop and extensions in the B-C loop (**fig. S10C**). Moreover, RedCAP FCP1 is the only antenna protein that interacts with PSI on the lumenal side, with the B-C loop protruding into a crevasse formed by PsaB, PsaI and PsaL in a “hand-in-glove” interaction. This creates a conserved, extended interface (**Fig. 3G-I**). Although RedCAP contains three family-specific chlorophyll-binding sites (**table S4**), this subunit forms the weakest EET pathways to PSI, with only one pair of chlorophyll molecules closer than 15 Å (**Fig. 3I**). Thus, RedCAP forms the most extensive FCP:PSI interactions, but the weakest expected EET transfers across the red lineage except for dinoflagellates (*27*).

In dinoflagellates, a RedCAP-to-Lhcf transition results in stronger EET interactions with PSI (**Fig. 3J-L**). Similar to the FCP3 rotation upon the loss of PsaO, we posit that this transition was enabled by the loss of the RedCAP gene in dinoflagellates (*39*). Here, the RedCAP loss broke the extended, contingent protein:protein interactions between FCP1 and PSI, allowing the FCP1 position to be filled by an Lhcf subunit. The main impact of the family change are the new chromophore interactions of Lhcf *versus* RedCAP. Rather than the limited EET interactions of RedCAP-FCP1 (**Fig. 3K**), Lhcf-FCP1 contains four chlorophyll molecules that were determined as key transfer positions from FCP1 to PsaB and PsaL (*27*) (**Fig. 3L**). Additionally, similar to Cg9-l FCP3, the dinoflagellate Lhcf-FCP1 is rotated ∼180° relative to the “standard” orientation, using helix A rather than helix C to interface with PSI (*26*, *27*) (**Fig. 3G,J**). This results in new protein:protein interactions, mainly to helix C of FCP11, and an increased distance to PSI (**Fig. 3J**).

### M. pyrifera’s second belt: likely Lhcf majority

In our structure, *M. pyrifera*’s PSI second belt is formed by FCP13/15/16/17/19/A/B (**Fig. 1**). Whereas FCP13/17 were assigned genes from the Lhcr and Lhcf subfamilies, respectively, the resolution of the remaining second-belt FCPs was insufficient resolution to assign protein identities. Thus, the FCP 15/16/19/A/B models, as well as FCP2 in the first belt, remained poly-alanine. We used map-model correlation scores (Q score) to identify the best-matching family for the poly-alanine models. For instance, Lhcf proteins have a shorter and straighter helix C than Lhcr and Lhcq proteins, providing useful features to differentiate families (**fig. S10**). We determined that Lhcf is the most likely subfamily for FCP15/16/19/A (**fig. S13**). The subfamilies of FCP2/B remained unassigned.

These results suggest that the majority of *M. pyrifera*’s second belt is composed of Lhcf proteins (FCP15/16/17/19/A), the exception being FCP13, which belongs to subfamily Lhcr. This Lhcf-majority composition would be unlike most other red-lineage PSI that contain a second belt. In diatoms and haptophytes, second-belt FCPs are predominantly Lhcq (*28–31*, *38*), whereas in cryptophytes the second belt is unanimously Lhcr (*22*, *23*) (**fig. S11**). However, dinoflagellates have a similar pattern to *M. pyrifera* in which all second belt FCPs are classified as subfamily Lhcf (*26*, *27*) (**fig. S12**). A shift from Lhcr to Lhcf in the second belt has EET implications, as the Lhcf subfamily lacks the Lhcr-specific chlorophyll-binding motifs.

### Chlorophyll losses/gains result in different EET pathways between *M. pyrifera* and diatoms

The red-lineage phylum Ochrophyta splits into two major clades, Chrysista and Diatomista, with vast molecular, morphological and metabolic diversity (*61*, *62*) (**fig. S1E**). For instance, whereas Diatomista is composed of unicellular organisms (e.g., diatoms), the Chrysista clade includes various types of multicellular macro- and micro-algae (*61*, *62*). Given this diversity, comparative studies of both clades are needed. To understand some of the photosynthetic differences between the Chrysista and Diatomista clades, we compared the chromophore composition of the PSI antenna between *M. pyrifera* and diatoms *Chaetoceros gracilis* and *Thalassiosira pseudonana* (*28–30*).

We focused on the FCPs that have been assigned in our structure, i.e., those with sufficient resolution to identify protein sequence and pigments (FCP 1/3/4/5/6/7/8/9/10/11/13), and their respective diatom analogues (**figs. S7, S11**). Overall, *M. pyrifera* contained fewer chromophore molecules per FCP than diatoms, with several conventional binding sites missing or unoccupied in the brown alga (*28–31*) (**tables S3-S4**). However, *M. pyrifera* also showed binding at locations not seen in diatoms or other red-lineage organisms. In diatoms, connections between FCP3/4/14, FCP10/11 and FCP7/8 have been determined as the strongest EET entry points into the PSI core (*28–31*). Below, we discuss chromophore differences and EET re-organization for *M. pyrifera* at these locations, together with their functional implications (**Fig. 4**).

**Fig. 4.**
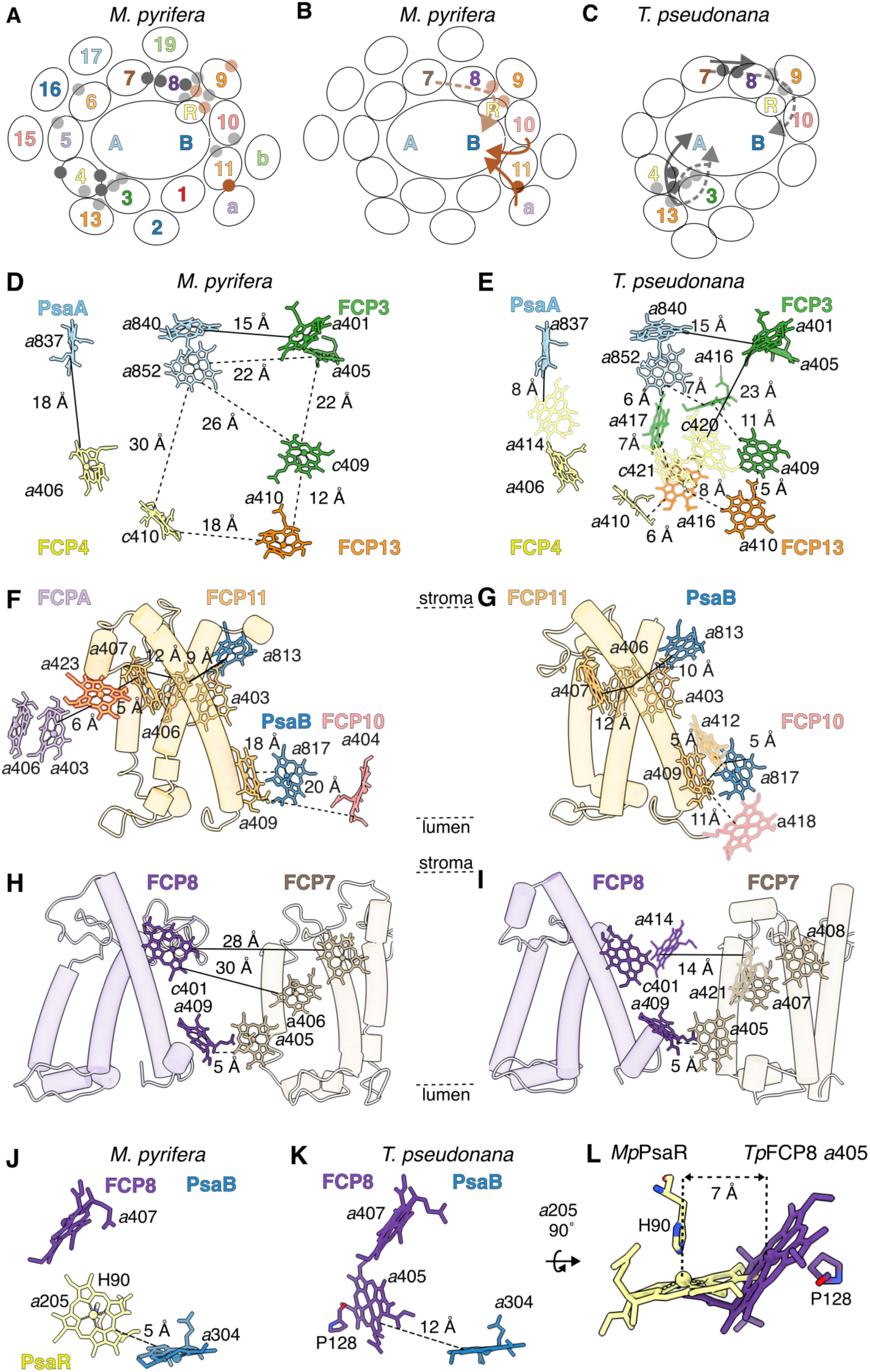
Key differences in PSI-FCP chromophore arrangement between *M. pyrifera* and diatoms. (**A-C**) Schematic summaries of chromophore arrangements in *M. pyrifera* (PDB: 9YGV, this work) (B) and diatom *Thalassiosira pseudonana* (PDB: 8ZEH) (*31*) (C). Dark grey, dark brown indicate chromophores on the stromal side; light grey, light brown, on the lumenal side. Subunits labeled without Psa/FCP prefixes for clarity. a, FCPA; b, FCPB. (A) Summary of chromophores lost (grey) or gained (brown) in *M. pyrifera* with respect to diatoms. (B-C) Key EET pathways in *M. pyrifera* (B) and diatom *T. pseudonana* (C). Stromal pathways in solid arrows; lumenal pathways in dashed arrows. Participating chromophores indicated with circles (dark, stromal; light, lumenal). (**D-L**) Key chlorophyll differences between *M. pyrifera* (D, F, H, J) and *T. pseudonana* (E, G, I, L) shown for various regions. Chromophores gained in *M. pyrifera* marked with thick brown border in *M. pyrifera* panels. Chromophores lost in *M. pyrifera* marked with grey border in *T. pseudonana* panels. Edge-to-edge distances between porphyrin rings indicated with solid lines for stromal connections and with dashed lines for lumenal connections. (D-E) Differences in FCP3/4/13-PsaA. (F-G) Differences in FCP11/A-PsaB. Note that FCPA is absent in the available diatom structures. (H-I) Differences in FCP7/8. (J-L) Chlorophyll migration from FCP8 (*T. pseudonana* chl *a*405) to PsaR (*M. pyrifera* chl *a*205). (L) Superposition of *T. pseudonana* (*Tp*) FCP8-chl *a*312 and *M. pyrifera* (*Mp*) chl *a*205, aligned by PsaR. Coordinating residues and approximate Mg-Mg distance between the chlorophyll molecules are shown.

### FCP3/13/4: chlorophyll losses suggest a weaker entry point via FCP3/4

*M. pyrifera* showed the loss of eight chlorophyll molecules in FCP3/4 relative to diatoms: five in the lumenal side and three in the stromal side (**Fig. 4A**). These losses increase the chlorophyll distance between FCP3/4 and PsaA with consequent decrease in EET efficiency and create a larger gap between FCP3/13 and PsaA, likely filled with lipids (**Fig. 4D,E**). The stromal losses of FCP4 also decrease the connectivity between FCPs 3/4/5, particularly due to the loss of the diatom-specific chl pair FCP4 chl *c*401-FCP4 *c*408 on the core-facing interface of FCP3/4 (**Fig. 4D,E).** Chlorophyll pairs encourage energy transfer towards them, acting as primary energy sinks and photo-protectants, significantly contributing to quantum efficiency. Thus, chlorophyll pairs are critical components of EET networks of large PSI supercomplexes (*63–65*). Overall, these changes imply that FCP3-4 are weaker EET entry points into PSI in *M. pyrifera* relative to diatoms, potentially a major functional difference between Diatomista and Chrysista ochrophytes. Given that the FCP3/4 chlorophyll complement of the other red-lineage organisms resemble *M. pyrifera* (*18–24*, *26*, *27*), it appears that the extensive FCP3/4/13 EET pathway is diatom-specific gain.

### FCP10/11: losses and gains suggest stronger predominance of stromal pathway

*M. pyrifera* showed two lumenal chlorophyll losses and one stromal gain in FCP10/11 (**Fig. 4F-G, fig. S14A)**. The lumenal changes increase the EET distances between FCP10/11, as well as between FCP11/PsaB from 11 Å to 20 Å and 5 Å to 18 Å respectively (**Fig. 4F,G)**. In contrast, the new stromal chl in *M. pyrifera* FCP11 (chl *a*423) appears to be ∼6 Å away from a chlorophyll *a* pair in FCPA (chl *a*403/*a406*), one of the new *M. pyrifera* antenna positions. Although the FCPA cryoEM map was reconstructed at lower resolution, it showed reasonable density for this chlorophyll pair (**fig. S14A**). All FCPs in our structure except for FCP1/3 contain a chlorophyll pair analogous to the chl *a*403/*a*406 pair proposed for FCPA, increasing the confidence for this assignment. The proximity of this FCPA chlorophyll pair and the new stromal chl *a*423 in FCP11 suggest that this is a new EET pathway from the second belt into PSI in *M. pyrifera* (**Fig. 4F,G**). Moreover, a fucoxanthin molecule (FCP11 Fx305) in the vicinity of these three chlorophylls likely provides further photoprotection and more efficient energy transfer (*25*) (**fig. S14A)**. A second FCP10/11 EET pathway into PSI exists on the stromal side *via* the FCP11’s Lhcr-specific chlorophyll (chl *c*415) and FCP10 chl *a*407. The ∼10 Å distance between these chlorophylls suggests they are also significant players in FCP10/11 transfer into the core. Replacement of FCP11 with a different FCP subfamily would increase the closest stromal distance with FCP10 to ∼23 Å. Overall, the lumenal losses, the stromal gain and the two stromal pathways in FCP10/11 imply a stronger predominance of the stromal transfer in this region *M. pyrifera* relative to diatoms, likely correlated to the new positions in the brown algal antenna (FCPA/B as well as additional unmodelled ones, **fig. S12**).

### FCP7/8: predominance of lumenal transfer into PsaR with a reconfigured EET pathway

In diatoms, the main FCP7/8 link is stromal (FCP7 chl *a*421 and FCP8 chl *a*414, ∼14 Å apart). Both of these stromal chlorophyll molecules are lost in *M. pyrifera*, resulting in an increased stromal distance between FCP7/8 of >28 Å (*30*) (**Fig. 4H-I)**. Additionally, stromal FCP8 chl *a*409 is also missing. Although the stromal pathway between FCP8-PsaR is maintained (**fig. S14B,C**), the chlorophyll losses imply a decreased EET transfer from FCP7/8 to PsaR on the stromal side in *M. pyrifera*. The lumenal side of *M. pyrifera* FCP8 also shows a chromophore loss (diatom FCP8 chl *a*405). However, this loss is compensated by a new chlorophyll in PsaR (chl *a*205 in our model), located ∼7 Å away (**Fig. 4J-L**). Thus, there is no *net* loss of lumenal pigments in FCP8. As discussed below, this reconfiguration implies a stronger lumenal entryway into the core at FCP7/8 in *M. pyrifera* relative to diatoms.

In diatoms, FCP8 chl *a*405 is coordinated by the mainchain carbonyl (and bridging water molecule) of Pro125 in helix C, ∼12 Å away from PsaB chl *a*833 (*28–31*). In *M. pyrifera*, the equivalent amino acid in that position (MpFCP8-Glu129) does *not* coordinate any chlorophyll molecules. Instead, the first transmembrane helix of PsaR has acquired His90, which now coordinates chl *a*205. This shift of chl *a*205 coordination from *Mp*FCP8 to *Mp*PsaR brings chl *a*205 closer to PsaB chl *a*835, now ∼5 Å away rather than ∼12 Å in diatoms (**Fig. 4J-L**). The CV/IHE amino acid motif that houses MpPsaR-His90 is conserved in the Phaeophyceae and various sister clades within Chrysista, but is missing in Diatomista ochrophytes other red-lineage phyla, suggesting it was acquired at the root of the Chrysista clade (**fig. S14D**). Although this replacement maintains the lumenal EET pathway from FCP8 into PsaB, it results in large changes in the chlorophyll coordination and surrounding protein environment relative to diatoms. In *M. pyrifera*, the chlorophyll molecule moves towards the interior of the complex and into the membrane (more hydrophobic), away from the lumen (more hydrophilic). Coordination type and environment are known to influence chl absorption and emission properties, as well as excited-state lifetimes (*66–70*). We thus predict significant differences in the light absorption and EET in this region of the PSI-FCP supercomplex in brown algae versus diatoms.

Additionally, diatoms’ FCP8 chl *a*405 is absent in red algae, cryptophytes and dinoflagellates but present in haptophytes, and the lumenal region of FCP8 also shows chromophore differences between diatoms (*28–31*). Together, these observations suggest that the FCP8 lumen is a hotspot for antenna diversification across the red lineage.

Overall, our analysis showed several differences in the chromophore arrangement and, consequently, expected EET pathways between *M. pyrifera* and the characterized diatoms (*28*–*31*). Spectroscopic studies of *M. pyrifera* and diatom PSI supercomplexes are needed to test the hypothesis that brown algae PSI has more prominent lumenal EET pathways and that the FCP8-PsaR entry into the core plays a larger role than the FCP13/3/4-PsaA entry (**Fig. 4**). Moreover, our cross-red-lineage analysis suggests that a prominent FCP3/4/13 EET pathway and the lack/presence of several chlorophyll molecules in FCP8 are specific to diatoms rather than to ochrophytes as a whole. Whether the differences we observed between *M. pyrifera* and diatoms are sustained across the Diatomista and Chrysista clades remains to be determined with further structural studies.

## Discussion

Here we present the structure of brown alga *M. pyrifera*’s PSI-FCP supercomplex (**Fig. 1**). PSI is well conserved with respect to the red lineage and other photosynthetic lineages (**fig. S8**). The FCP antenna in our structure contained two belts arranged differently from diatoms and other red-lineage organisms (**Fig. 1, fig. S11**). Preparations with different detergent conditions showed that larger antenna architectures exist for *M. pyrifera*’s PSI-FCP supercomplex (**Fig S12**). *M. pyrifera*’s FCP phylogeny supported a shared topology across the red lineage (*38*, *39*) (**fig. S9**). Our comparisons of structures and FCP subfamily membership across antennae of the red lineage suggested drivers of conservation and diversification at specific FCP positions, as discussed below (**Fig. 2-3, fig. S11-S15**). Our comparison of chromophore differences between *M. pyrifera* and diatoms determined a remodeling of EET pathways, with a weaker entry into PSI *via* FCP3/4, a stronger stromal path for FCP10/11 with additional links for the new second-belt positions, and a re-configured lumenal path via FCP7/8-PsaR in the brown alga (**Fig. 4, fig. S14-S15**). Collectively, our analyses identified differences between the Chrysista and Diatomista clades of ochrophytes and provide new insights into the evolutionary relationships across the red lineage (**fig. S1, S14-S15**).

### Drivers of FCP-subfamily occupancy in each antenna position

Based on our comparative structural and evolutionary analysis, we propose that contingent protein:protein interactions, as well as a previously unrecognized hydrophobic mismatch, are key drivers of the positional conservation of FCP subfamilies and PSI-FCP function.

#### Contingent FCP:PSI protein:protein interactions, released by gene loss

Similarities in sequence, subunit composition and architecture of the supercomplex are strongest in PSI, weakening with increasing distance from the core (**fig. S11**). Thus, the FCP first belt is more conserved in architecture and subfamily composition than the second, as seen in the conserved Lhcr majority across the red lineage (**fig. S11, table S5**). We posit that the conservation of the first belt is driven by the sequence conservation of the PSI subunits, to maintain the PSI:FCP binding interactions. The Lhcr first-belt majority is further aided by subfamily-specific features, e.g., Lhcrs’ aromatic residues that allow for FCP:FCP interactions not seen in other subfamilies (**Fig. 2**). Lhcr proteins are also well suited to the first belt thanks to their additional family-specific chromophore binding site (chl 415) (*38*) (**Fig. 2**, **fig. S10-S11**). The family-specific chlorophyll may be particularly important to allow FCP10/11 to efficiently receive excitation energy from the new *M. pyrifera* positions in the second belt and transfer it to PSI (**Fig. 4F-G**).

Another example of conserved protein:protein interactions with the core driving positional conservation of families is FCP1 (**fig. S11**). FCP1 is conserved as a RedCAP protein in red algae, cryptophytes, ochrophytes and haptophytes despite a weak EET connectivity and the presence of multiple FCP subfamilies with additional chromophores in the latter two clades (*39*) (**fig. S11**). This strongly suggests that the FCP1 conservation is driven by the contingent binding surface rather than by RedCAP’s functional significance for light absorption or EET (*38*) (**Fig. 3K-L**).

The historical contingency of binding interactions can be broken by gene loss, which allows the “empty” position to be filled in new ways. We propose that this explains the subfamily switch and rotation for FCP1 and FCP3 (**Fig. 3, fig. S11**). For FCP1, loss of the RedCAP gene in dinoflagellates led to a replacement of the FCP1 position with a pre-existing Lhcf protein (*38*). A similar situation is seen with FCP3, which switched from Lhcr to CgLhcr9-like in haptophytes, ochrophytes and dinoflagellates after the loss of binding partner PsaO (nuclear-encoded core subunit in red algae and cryptophytes). In both cases, the switch and rotation allowed for new protein:protein interactions and new EET networks to emerge (**Fig. 3**).

#### Hydrophobic mismatch

When the length of the transmembrane portions of proteins does not match the thickness of the surrounding membrane, there is a “hydrophobic mismatch” with significant energetic costs (*71*). Proteins and membranes can undergo structural changes to minimize the penalty. For instance, the membrane may stretch, compress, distort or switch the type of lipids to alter its thickness (*51–55*). Proteins may change the tilt or bend of their transmembrane helices to fit within the existing membrane, which may alter protein:protein interactions and lead to sorting into lipid microdomains in the membrane (*51–57*). Hydrophobic mismatch is exploited in at least two photosynthetic membranes from bacteria to the green lineage: i) protein translocation by the Tat translocon (*72–74*) and ii) PSII activity during the energy-dependent component (qE) of non-photochemical quenching (NPQ) in the green lineage (*75*, *76*). In the latter, high light leads to lipid reorganization (particularly, of digalactosyldiacylglycerol, DGDG), leading to membrane thinning from ∼40 Å to ∼30 Å (*75*, *76*). The consequent mismatch imposed on PSII leads to the tilting of the LHC helices, resulting in “flattened” PSI-LHCII supercomplexes. These flat supercomplexes seed further membrane thinning, as well as supercomplex aggregation and sequestration into specific membrane microdomains (*75*, *76*, *56*). Whereas *M. pyrifera* does not possess the qE component of NPQ (*44*, *45*), DGDG is a major component of brown algal chloroplast membranes (*77*, *78*). Additionally, Tat translocon genes have been predicted in brown algal model *Ectocarpus siliculosus* (*79*), suggesting that hydrophobic mismatch is at play in brown algal chloroplasts.

In *M. pyrifera*’s PSI-FCPs, we observed significant differences in the length of the C-helix across FCPs (∼40 Å to ∼24 Å at its largest, FCP5 and FCP17, respectively) and across families (average of ∼31.5 Å for Lhcr vs ∼23.2 Å for Lhcf, **Supplementary Data File S2**). These differences are in the range of values established to be significant for mismatch-mediated PSII regulation in plants (*75*, *76*). Given that the Lhcr subfamily shows the longest C-helices and a conserved first-belt majority (**Fig. 2F-J**), we hypothesize that hydrophobic mismatch is a key driver of FCP subfamily occupancy and of Lhcr maintenance in the first belt across the red lineage. Minimization of mismatch with the membrane thickness around PSI would favor the incorporation of FCPs with the thickest helices in the first belt, i.e., Lhcr proteins.

Moreover, we hypothesize that hydrophobic mismatch is a regulatory mechanism of PSI function in brown algae and the red lineage more broadly (**Fig. 2**). For instance, similar to green-lineage PSII, high light could lead to the rearrangement of DGDG in the brown algal thylakoid. The consequent membrane thinning could lead to a conformational change in the C-helices’ tilt or curvature, leading to the destabilization of PSI:FCP or FCP:FCP interactions that could alter PSI’s photosynthetic activity in high light. DGDG levels have been observed to decrease under high light in *Undaria pinnatifida* (wakame kelp), leading to changes in chloroplast morphometry and photosynthetic efficiency (*78*). A role for hydrophobic mismatch in regulating PSI function under high light could compensate for the lack of qE-based photoprotection through PSII in *M. pyrifera* and brown algae (*44*, *45*, *80*, *81*).

In parallel, the differences in the length of transmembrane helices across the PSI-FCP supercomplex imply localized differences in membrane thickness (**Fig. 2F-K**). It is understood that membrane thickness affects the dielectric properties of the membrane, leading to differences in the electric field across the membrane (*47*, *54*). Moreover, changes in the electric field affect the energetics and kinetics of the charge separation reactions of the photosystems (*48*) as well as the absorption spectrum of chromophores (*49*, *50*). Thus, we hypothesize that, by creating different membrane thicknesses, the length of the C-helix of FCP proteins allows for the “tailoring” of the local electric field, fine-tuning the local electric environment, and thus absorption properties, of different regions of the antenna. We hypothesize that this provides an additional selective advantage for Lhcr proteins at particular locations of the first belt. Moreover, the minimization of hydrophobic mismatch could also explain why FCP rotations are favored in the first belt upon gene loss and subfamily switches (**Fig. 3**). Biophysical experiments with membrane-reconstituted PSI supercomplexes will shed light on this hypothesis.

### Diversification of the second belt

Most subfamily and chromophore diversification seen across red-lineage organisms occurs in the second and further belts (**fig. S11**). Some subfamily changes, e.g., FCP9 changing from an ancestral Lhcr to Lhcf and FCP13 changing from Lhcr to Lhcf or Lhcq, follow the loss of ancestral chromophore-binding motifs (e.g., Lhcr’s N-terminal SxS/AL/IP that binds family-specific chl 415, **fig. S10**, **table S3-S5**). However, relative to the “most ancestral” state available, many of these changes appear class- or species-specific, or without a clear evolutionary pattern (**fig. S11, table S5**). The second-belt majority is also inconsistent between red-lineage phyla (Lhcr in cryptophytes, Lhcq in haptophytes, Lhcf in dinoflagellates and likely in *M. pyrifera*), and even between diatoms (Lhcr/Lhcf in *T. pseudonana*; Lhcq in *C. gracilis*) (*18–31*) (**table S5**).

The diversification of the outer-belt composition has been explained as an adaptation to clade-specific environments, enabled by organism-specific expansions of FCP subfamilies such as Lhcf, Lhcq and Lhcx (*6*, *38*, *39*, *18*). We propose that the theory of constructive neutral evolution (CNE) provides an alternative to understand this diversification (*16*, *82*). CNE posits that complexity can arise neutrally rather than by positive selection. Key components of the CNE framework are excess capacity (e.g., “gratuitous” components such as from gene duplications and family expansions), epistasis (context-dependent phenotypic effects of mutations, e.g., interdependencies due to protein:protein interactions) and biased variation (e.g., the fact that mutations that reduce function such as the loss of a chromophore-binding motif are more common than those conferring enhanced function), as well as random genetic drift and purifying selection (*16*, *82*). In the case of the PSI antenna, the FCP duplication and expansion from Lhcr into Lhcf, Lhcq, Lhcx, Lhcz and Cg9-l subfamilies would provide excess capacity (*38*). For instance, we determined that *M. pyrifera* contains 22 Lhcf, 6 Lhcq, 8 Lhcx, in addition to 17 Lhcr, 2 RedCAP, 1 Lhcz and 1 Cg9-l genes (**fig. S9**). Many of the new genes might contain slightly deleterious changes, e.g., one fewer chromophore in the lumen as is the case for Lhcf *vs* Lhcr. The plentiful FCP capacity would lead to many gratuitous interactions between FCPs while assembling the outer belts, e.g., incorporating an Lhcf rather than Lhcr protein in the FCP17 position, as is the case between *M. pyrifera* and cryptophytes (**fig. S11**). The suboptimal mutation of the Lhcf might be rendered neutral by epistasis, e.g., if small mutations in its Lhcr partner result in a favorable binding interaction. Then, this new composition might drift to fixation, effectively “locking in” the originally gratuitous interaction, *even if* the original change (incorporating an Lhcf) was slightly deleterious on its own. Thus, the interdependency leads to an increase in complexity that arose neutrally and not necessarily by positive selection (*16*, *82*). We posit that constructive, neutral processes may be shaping the antenna architecture and composition, particularly at positions that do not play critical roles in the central EET pathways. We emphasize that we are *not* proposing this to be the only evolutionary process at play in the evolution of PSI-FCP supercomplexes.

The PSI core is a highly efficient photochemical machine achieving ∼99% quantum efficiency, surpassing PSII (∼85%) (*83*, *63*, *65*). It has been previously proposed that this high efficiency allows for the expansion of the antenna size, as PSI can accommodate increased travel distances and times from the outer belts without a too-detrimental compromise to the quantum efficiency (*63*, *65*). For instance, the haptophyte *Emiliania huxleyi* achieves 95% efficiency with its eight-belt FCP antenna (*25*). In terms of CNE, the high PSI efficiency would provide a permissive landscape to incorporate the FCP excess capacity, leading to larger, diversified, more complex antennae that may not necessarily increase fitness. Thus, the diversification of PSI’s outer antenna across the red lineage need not be positively adaptive; instead, it may be a response to neutral processes enabled by the permissivity of PSI’s high efficiency.

This hypothesis can be tested with biophysical studies of natively purified PSI reconstituted with different FCP combinations (*84*). It has also been recognized that comparative structural analyses of multi-subunit complexes across related organisms, such as the one presented here, is a helpful approach to infer the functional impact of increases in complexity (*16*, *17*). Thus, structural and functional comparisons across additional key red-lineage organisms will reveal whether the diversification of PSI-FCP supercomplexes is the result of positive selection or neutral constructive evolution.

### Implications for the endosymbiotic origin(s) of the red lineage

The endosymbiotic relationships between the phyla of the red lineage remain unclear, with three models consistent with recent molecular-timescale-constrained phylogenetic analyses (*14*, *15*) (**figs. S1A-D**, **S15A-C**). A model from Stiller *et al*. posits serial endosymbioses of red algae to cryptophytes to ochrophytes to the ancestor of haptophytes and myzozoa (dinoflagellates) (*85*) (**fig. S15A)**. In contrast, Bodyl *et al*. propose serial endosymbioses from red algae to cryptophytes to the ancestor of ochrophytes and haptophytes, followed by an engulfment of a haptophyte to give rise to myzozoa (*86*) (**fig. S15B**). Additionally, Pietluch *et al*. propose that there were two separate secondary endosymbioses of red algae followed by separate tertiary endosymbioses from cryptophyte to haptophyte and from ochrophyte to myzozoa (*15*) (**fig. S15C**). Comparing the structures and composition of photosystems—assemblies of plastid- and nuclear-encoded subunits—provides a complementary approach to evaluate evolutionary relationships (*17*). Given the different roles of ochrophytes in the three models, examining Chrysista and Diatomista structures is particularly helpful.

Clear support for the Pietluch model comes from FCP9/13, where cryptophytes and haptophytes are closely aligned, but different from ochrophytes and dinoflagellates, themselves closely aligned (*15*) (**fig. S15C**). These two Lhcr subunits lack the otherwise Lhcr-specific chl 415 in *M. pyrifera*. The main driver of the chl loss is the fact that their N-terminal loop adopts a conformation unlike that seen in the rest of the Lhcr subunits. In *M. pyrifera*’s FCP9/13, the N-ter loop travels away from the core rather than forming a core-facing loop with the chlorophyll-binding turn (**fig. S15D**). An analogous situation in seen in diatoms and dinoflagellates (**fig. S15E**). In the latter, dinoflagellate FCP9/13 belong to the Lhcf subfamily, with a short 15-amino-acid helix C, but otherwise fully align with *M. pyrifera*’s FCP9/13 structure. In contrast, FCP9/13 contain the traditional Lhcr N-terminal motif and structure in red algae, cryptophytes and haptophytes (**Fig S15E**).

The pattern of inheritance and protein arrangement in the PsaK/O-FCP2/3 region provide additional cases (**fig. S15F-G**). PsaO is encoded in the nucleus of red algae and of cryptophytes and is absent in the other red phyla (*58*, *59*, *24*, *26–30*). The pattern suggests that red algal PsaO was transferred to the nucleus of the ancestral cryptophyte upon secondary endosymbiosis and was lost in subsequent endosymbioses due to failure to transfer to the nucleus from the engulfed nucleomorph. This implies PsaO’s nuclear-transfer failure happened once (cryptophyte to ochrophyte, **fig. S15A,3**) for the Stiller model, twice for the Bodyl model (cryptophyte to ochrophyte, **fig. S15B,3i**; cryptophyte to haptophyte, **fig. S15B,3ii**) and twice for the Pietluch model (cryptophyte to ochrophyte, **fig. S15C,3i**; red alga to ochrophyte, **fig. S15C,2i**). The loss of PsaO from PSI is perfectly correlated with the subfamily switch and physical rotation of FCP3 (**Fig. 3**, **fig. S11**). The facts that ochrophyte, haptophyte and dinoflagellate show the same new FCP3 subfamily (Cg9-l) and high similarity in FCP3 structure, orientation and PSI-binding, and that these clades are in stark contrast to red algae and cryptophytes, is most parsimoniously explained by the Stiller model, which posits a common ancestor the ochrophyte, haptophyte and dinoflagellate clades (*85*) (**Fig. 3A-B**, **figs. S11**, **S15B**). The filling of the PsaO gap with the unusual FCP3 family and orientation in two independent events, as expected from the Bodyl and Pietluch models would be less likely (*15*, *86*). A single nuclear-transfer failure would suffice for the Bodyl model if the engulfed cryptophytes for both ochrophytes and haptophytes belonged to a photosynthetic clade already lacking PsaO. The absence of reported extant photosynthetic cryptophytes lacking PsaO decreases the likelihood of this scenario, although we note that some cryptophytes do not assemble PsaO into PSI in certain growth conditions (*23*). The second case of subfamily and rotational switch, i.e., FCP1’s RedCAP-to-Lhcf change, does not provide evidence to disambiguate the three models, as the modification occurred in dinoflagellates, i.e., upon the terminal endosymbiosis in all models (**Fig. 3G-L**, **fig. S15A-C**,**H-I**).

Lastly, PsaK is encoded in the plastid of red algae, cryptophytes and in the nucleus of haptophytes (*58*, *59*, *87*). This is consistent with a nuclear transfer from the cryptophyte nucleomorph to the haptophyte nucleus in the Bodyl and Pietluch models (**figs. S15B,3ii**; **S15C,3i**), with a subsequent nuclear-transfer failure from haptophytes to dinoflagellates (**fig. S15B,4**) in the Bodyl model. In the Stiller model, the PsaK pattern implies a nuclear-transfer failure from cryptophyte to ochrophyte (**fig. S15A, 3i**) and then re-gain from ochrophyte to haptophyte (**fig. S15A,4i**), or other non-parsimonious scenarios that decrease support for this model (*85*). Additionally, FCP sub-family changes in FCP2 in the vicinity of PsaK/O are also more consistent with the Bodyl model: FCP2 is an Lhcr protein in cryptophytes, haptophytes and dinoflagellates, whereas it is Lhcq in ochrophytes (*18–31*) (**figs. S11, S15F-G**).

These cases illustrate how structural comparisons of composition, conformations and interactions can complement and shed light on phylogenetic hypotheses. Any accepted evolutionary model will need to be consistent with the protein and chromophore features revealed by high-resolution structures of PSI, PSII and other multi-subunit protein assemblies (*18–31*). Structures from additional red-lineage clades, coupled with a better understanding of the evolutionary history of the specific FCPs found in the structures and of the inferred ancestral organisms that were engulfed, are needed to elucidate the relationships in the red lineage.

### Outlook

Our structural characterization and comparative analysis of *M. pyrifera* PSI-FCP allowed us to identify i) Chrysista- and Diatomista-specific features with functional implications on EET, ii) drivers for antenna conservation in the first belt, including a potential new regulatory process by hydrophobic mismatch, iii) the feasibility of a constructive, neutral evolutionary framework for the diversification of the outer belts and iv) cases for support for and disagreement with three evolutionary models for red-lineage evolution. Further structural comparisons across the red lineage, as well as biochemical, biophysical and genetic analyses will test our hypotheses. Moreover, this first cryoEM structure of a brown algal complex lays the groundwork to understand kelp’s high photosynthetic productivity “from the bottom up”, providing a biochemical basis for the certification of kelp forests as blue carbon ecosystems (*34*).

## Materials and Methods

### Chloroplast purification

Fresh *Macrocystis pyrifera* blades were obtained from Monterey Bay Seaweeds. Their chloroplasts were isolated based on previous protocols, with modifications (*88–92*). For each isolation preparation, ∼1 kg of fresh fronds was rinsed with deionized water and treated with H₂O-HCl (pH 6.0) for 30 min to remove surface-adsorbed materials. The tissue was then rinsed, finely chopped and incubated twice in a 0.1 M sodium citrate buffer (pH 8.0) at 4 °C under gentle stirring to remove alginates (*93*). Samples were centrifuged at 5,000 x g for 10 min and washed with deionized water to remove residual citrate. The pellet was resuspended in ∼1:8 g:mL of homogenization buffer (50 mM Tris, 150 mM NaCl, 10 mM phosphate, 1 mM EGTA, 0.35 M mannitol, 1% w:v PVP-40, 0.1% w:v BSA, 8 mM cysteine and boric acid at pH 8.0). To further decrease the viscosity of the resuspension and aid in organelle isolation, recombinant alginate lyase (Alg2A from *Flavobacterium* sp. S20, 6 mg L⁻¹) (*94*) was added, and the mixture was incubated for 1 h at 4 °C. The tissue was then homogenized using a 4-L blender (Waring, Stamford, CT, USA, model: CB15BU) at high speed for three consecutive 5-second cycles, filtered through two layers of Miracloth (MilliporeSigma, Burlington, MA, USA) and centrifuged at 5,000 x g for 45 min at 4 °C. The chloroplast pellet, typically yielding ∼250 g kg⁻¹ of initial tissue, was washed in a buffer containing 50 mM HEPES, 0.3 M mannitol, and 1 mM EDTA (pH 7.5), and further purified by layering it on top of a 2-step Percoll gradient (40% v:v over 80% v:v) (Sigma-Aldrich, St. Louis, MO, USA) diluted in a 20 mM HEPES buffer at containing 0.33 M sucrose at pH 7.5. The gradient was centrifuged at 3,000 x g for 20 min at 4 °C (*95*). The chloroplast-enriched fraction was collected manually, washed twice with a buffer containing 20 mM HEPES, 1 mM EDTA, 50 mM NaCl, and 10% glycerol, adjusted to pH 7.5 with NaOH, by centrifuging at 4,000 x g for 5 min. The pellets were then aliquoted in volumes of 2 mL, and flash-frozen in liquid nitrogen and stored at –70 °C.

### Chloroplast membrane wash

To isolate thylakoid membranes, the purified chloroplasts were subjected to a hypotonic shock by resuspension in chilled Milli-Q water at a ratio of 10 mL:1 g water:membranes and homogenized with 100 strokes using a glass homogenizer with a tight pestle. Subsequently, 3M KCl was added to adjust the suspension to a final concentration of 150 mM KCl and re-homogenized with an additional 100 strokes. Membranes were collected by centrifugation at 45,000 x g. The pellet (membrane fraction) was solubilized in buffer (20 mM Tris-HCl, 150 mM NaCl, 1 mM EDTA, 10% v:v glycerol, 10 U mL⁻¹ DNase I, 2 mM DTT and 0.002% PMSF, at pH 7.4) and homogenized with another 100 strokes using a glass homogenizer and tight pestle. After final homogenization and centrifugation at 45,000 x g for 30 min at 4 °C, the membrane fraction was obtained as a brown pellet enriched in thylakoid membranes. The chlorophyll concentration of the sample was determined spectrophotometrically following the equations of Jeffrey and Humphrey (*96*). To extract the chlorophyll, samples were incubated with 0.1% Triton X-100 (Thomas Scientific, Swedesboro, NJ, USA) for 10 min at room temperature, followed by dilution in 90% acetone and incubation with gentle tumbling for 10 minutes at room temperature. Samples were then spun at 14,000 x g for 15 minutes and the supernatant was collected. The absorption of the supernatant was measured from 350 nm to 750 nm at 2 nm intervals, using a Spectramax M2spectrophotometer (Molecular Devices, San Jose, CA, USA). Membranes were aliquoted in a volume of 1 mL at 0.7 mg mL^-1^ of chlorophyll content, flash-frozen in liquid nitrogen and stored at –70 °C in a buffer containing 20 mM Tris, 50 mM NaCl, 1 mM EDTA, 2 mM DTT, 0,002% PMSF and 30% glycerol at pH 7.4.

### PSI-FCP supercomplex extraction and purification

To extract PSI-FCP complexes, the washed membranes were solubilized in 1% w:v digitonin (MilliporeSigma, Burlington, MA, USA) (detergent-to-chl w:w ratio of 73:1, or 12:1 for dataset in **fig. S12**) using extraction buffer (15 mM HEPES, 20 mM KCl, and 5 mM NaCl at pH 7.5) for 1 h at 4 °C under gentle tumbling. The solubilized material was clarified by centrifugation at 16,000 x g for 15 min, and the supernatant was concentrated to 100–200 µL using an Amicon Ultra Centrifugal Filter, 100 kDa MWCO (MilliporeSigma, Burlington, MA, USA). Linear sucrose gradients (20–50% w:v) were prepared in extraction buffer (15 mM HEPES pH 7.5, 20 mM KCl, and 5 mM NaCl) containing 0.01% GDN (Anatrace, Maumee, OH, USA). The concentrated sample was layered onto the gradients and ultracentrifuged at 263,600 x g for 22 h at 4 °C. Gradients were fractionated using a Gradient Station (Biocomp Instruments, Fredericton, NB, Canada). Fractions with a maximum absorbance peak at 676–678 nm were collected and selected for cryoEM sample preparation below.

### HPLC pigment analysis

Pigments were extracted from *M. pyrifera* PSI-FCP using acetone with 0.1% Triton X-100 v:v. Pigment analysis was performed by high performance liquid chromatography (HPLC) on an Agilent 1100 Series system equipped with a diode array detector and an Agilent Eclipse XDB column (5 μm, 4.6 x 150 mm). Separations were carried out at 20 °C with a flow rate of 1 mL min^-1^ using solvent A (MeOH/H_2_O = 90:10 [v:v]) and solvent B (ethyl acetate). The gradient was programmed as follows: 0-20 min, 0-100% B; 20-22 min, 100% B; 22-23 min, 100-0% B; and 23-28 min, 0% B. Pigments were detected at 445 nm with fill spectra collected from 300-800 nm, and identified by comparison to authentic standards of chlorophyll *a*, *c*, β-carotene, fucoxanthin, violaxanthin (Sigma-Aldrich, St. Louis, MO, USA) and zeaxanthin (AmBeed, Buffalo Grove, IL, USA). Quantitative standard curves were prepared in 100% acetone using the following concentrations: 0.5 μg mL^-1^, 1 μg mL^-1^, 5 μg mL^-1^, 10 μg mL^-1^, 50 μg mL^-1^, and 100 μg mL^-1^. Absolute quantification of PSI pigments was achieved by measuring peak areas. Carotenoid standards (zeaxanthin, violaxanthin, and fucoxanthin) contained isomeric mixtures and peak areas were summed for quantification.

### CryoEM grid preparation and data collection

The sample used for cryoEM grid preparation consisted of the above sucrose-gradient-purified PSI-FCP supercomplex from *M. pyrifera* at a final chlorophyll concentration of 1.0 mg mL⁻¹. Selected fractions from the purification above was buffer-exchanged into a buffer containing 15 mM HEPES (pH 7.5), 20 mM KCl, 5 mM NaCl, 0.01% (w:v) GDN (Anatrace, Maumee, OH, USA), and 0.1% (w:v) digitonin (MilliporeSigma, Burlington, MA, USA). CryoEM grids (Quantifoil 1.2/1.3 300 mesh Gold, GmbH, Germany) were glow-discharged for 30 s at 30 mA (PELCO easiGlow, Ted Pella Inc, Redding, CA, USA), followed by a 10 s hold prior to sample application. Grids were prepared by double blotting procedure. First, 2 µL of sample were applied, held for 30 s and manually blotted for 5 s outside the plunge freezer. Subsequently, 4 µL of sample were added, incubated for 10 s, and blotted for 8 s under controlled conditions (18 °C and 90% relative humidity) in a Leica EM GP2 cryo-plunger (Leica, Wetzlar, Germany). Grids were then plunge-frozen in liquid ethane using the same instrument.

CryoEM data were collected using a 300 kV microscope equipped with a Falcon 4i detector operated in electron counting (EC) mode at SLAC National Accelerator Laboratory, Stanford, CA, USA. Data acquisition was performed using EPU software. A total dose of 50 e⁻ Å⁻² was fractionated into 40 frames over an exposure time of 4.95 s, resulting in a dose rate of 1.25 e⁻ Å⁻² frame⁻¹. Movies were collected at a physical pixel size of 1.217 Å, corresponding to a magnification yielding 1.27 Å pixel⁻¹ on the detector. A total of 15,690 micrographs were acquired, with a fluence of 6.82 e⁻ pix⁻¹ (10.10 e⁻ Å⁻² s⁻¹).

### CryoEM image processing and composite map generation

Motion correction and contrast transfer function (CTF) estimation were performed using CTFFIND4.1 (*97*) in Relion v4.0.1 (*98*). Particle picking was initially guided by manual picking and subsequently used to train a Topaz model (*99*). A total of ∼3 million particles was extracted at a pixel size of 4.868 Å/pixel and transferred to cryoSPARC v4.4.1 (*100*). After successive rounds of 2-dimensional classifications with removal of undesirable classes, 426,437 particles were retained. Three *ab-initio* volumes were generated and used as inputs for heterogeneous refinement, yielding a dominant class of 164,731 particles (38%). The 164,731-particle stack was re-extracted at the original pixel size of 1.217 Å pixel⁻¹ in RELION v4.0.1 and re-imported into cryoSPARC. Non-uniform refinement (NU-refine) was performed (*101*), followed by core-focused Local Refinement (with “estimate per-particle scale” enabled) using a soft binary mask around the PSI– core generated in UCSF ChimeraX v1.10.1 (*102*). Particles clustering in a minor peak of a bimodal distribution of per-particle scale factors (value < 0.65) were discarded, resulting in a final data set of 133,000 particles.

After the initial refinement of the PSI core from 133,000 particles, the same particle set was used to obtain reconstructions of the antenna proteins. For the FCP subunits in the first belt, two complementary strategies were employed depending on the occupancy of each subunit. For FCP1, FCP3, FCP7, FCP8 and FCP10, where the density was stable and well-defined, individual soft masks were generated in ChimeraX around each subunit and imported into cryoSPARC. Each mask was created with a soft padding of six pixels with no dilation, and subsequently used for local refinement of the corresponding FCP. The other first belt FCP subunits (FCP2/4/5/6/9/11) required an additional classification step to separate particles in which the FCP density was not strongly present in the definitive core map. Subsequently, a masked classification using a soft mask of the corresponding FCP density was performed for each FCP. This was followed by local refinements focused on each FCP using specific soft masks. Refinement of the second-belt FCPs required a hierarchical alignment approach due to their peripheral and more flexible association with the PSI core. For the refinement of each second-belt FCP, particle stacks were first aligned using the neighboring first-belt subunit as a reference. Specifically, FCP3 was used to align FCP13; FCP5 for FCP15; FCP6 for FCP16; FCP7 for FCP17; FCP19, FCP1 for FCPA; FCP11 for FCPB.

After these alignments by the first belt, soft masks of each second-belt FCP density were applied around each second-belt subunit to perform masked classification into “present” and “absent” classes. Local refinements using the same second belt masks were subsequently carried out on the “present” subsets. Maps originating from the local refinement of the PSI core and of the individual FCPs were aligned to the overall map of the PSI-FCP supercomplex (before focused refinements) in ChimeraX and integrated into a composite map using phenix.combine_focused_maps in Phenix (*103*).

### Model building

Sequences of *M. pyrifera*’s PSI (core subunits) were identified by BLAST searches (*104*) using the annotated subunits of *Ectocarpus siliculosus* (Genome ID: CAID01000000) as queries against the Macpyr2 genome (*36*) (*Macrocystis pyrifera* CI_03 v1.0) in the Phycosm database (*37*). The top hits selected for each subunit were used to generate initial models with AlphaFold 3 (*105*). These structures were rigid-body fit into cryoEM density maps in ChimeraX and manually real-space-refined in Coot 0.9.8.96 (*106*). The PsaR subunit was not initially detected by BLAST searches. Instead, its density was identified in the cryo-EM map, and the corresponding region was carved, resulting in a map of 2.6 Å resolution. The map was subjected to ModelAngelo (*93*) to obtain a putative sequence assignment. The sequence retrieved by ModelAngelo was then used as a BLASTP query against the NCBI ClusteredNR dataset in GenBank (*107*), yielding a *M. pyrifera* homologue as the top hit. A structural prediction model was generated with AlphaFold3, fit into the density and validated through detailed visual inspection of side-chain densities.

To begin the assignment of the protein identities of the FCP densities in our maps, we reasoned there was a high likelihood that many of the *M. pyrifera* FCPs would be the homologues of the identified *C. gracilis* sequences(*29*). Therefore, we started by searching for the top hits of each *C. gracilis* FCP sequence in the *M. pyrifera* genome in PhycoCosm (*36*) using BLASTP searches (E-value threshold 1.0 × 10⁻⁵ with BLOSUM62; relaxed to 1.0 × 10⁻³ where necessary). The three top-scoring *M. pyrifera* hits for each diatom FCP query were retained as candidates. Structural predictions for each top hit, including seven chlorophyll molecules to stabilize the fold, were generated with AlphaFold3 (*105*). The predicted structures were rigid-body fit into the cryoEM density in ChimeraX (*102*) and manually refined in Coot (*106*) with particular attention to helical register assignment, bulky side chains and canonical chlorophyll-binding motifs. To determine the true protein identity among the three top candidates, the sequences of the three top hits were aligned, and the positions showing the highest amino-acid structural variability were noted (e.g., a position with a bulky residue in one candidate and a small residue in the other candidates). The fit of the predicted model of each candidate to the map was manually inspected, and sequences were compared to each other in detail, with particular attention to the positions of major differences. This assessment of the compatibility of each candidate with the density features led to an unambiguous assignment for the majority of first-belt FCPs (FCP1–FCP11, excluding FCP7) and second-belt subunits where the resolution was sufficient for molecular assignment (FCP13 and FCP17). FCP7 (first belt) required a distinct strategy, as BLASTP searches with the diatom and *Ectocarpus* queries yielded ambiguous candidates. Thus, we took a different approach, similar to the above for PsaR. The cryo-EM density corresponding to FCP7 was subjected to ModelAngelo (*93*), which provided a putative sequence. This sequence was subsequently used as a query in a tBLASTn search against *M. pyrifera* transcriptomic database in Phycosm (*36*). The best hit corresponded to an unannotated transcript, from which the open reading frame was translated into a protein sequence. AlphaFold3 models of the top hits were generated, fit, and assessed, leading to the unambiguous assignment of FCP7 according to the same criteria applied for the other FCPs. The second-belt FCPs whose resolution was not sufficient to assign a protein sequence (FCP15/16/19/A/B) were modeled as a poly-alanine chain, based on the FCP17 model that was then manually fit into the density as best as possible.

Ligand assignment was guided by homologous diatom PSI-FCP supercomplex structures (*108*–*110*) and refined by visual inspection of the density with manual adjustments when required. Chlorophyll molecules were modeled where porphyrin-like densities were clearly resolved. chl *a* and chl *c* were distinguished based on the continuity of phytol chain density and based on previous assignment in the highest-resolution diatom structure available (*108–110*). The atoms of the chl *a* phytol chain were removed where the density did not allow for atomic modeling. Carotenoid placement was initially guided by the positions observed in homologous diatom and related PSI-FCP structures (*108–110*). The assignments were then refined by visual inspection of each model, with fucoxanthin distinguished from violaxanthin based on the shape of the head group.

Following the modeling of each FCP subunit and ligand, the complete PSI-FCP supercomplex was assembled around the PSI core in ChimeraX. The integrated model was then subjected to real-space refined against the composite cryo-EM map in Phenix (*103*, *111*), applying secondary structure and geometry ligand restraints. Final validation was performed with MolProbity (*112*) in Phenix.

### FCP molecular phylogeny reconstruction

The full set of FCP genes in *Macrocystis pyrifera* were identified from the Macpyr2 (CI_03 v1.0) genome resource (*36*) in PhycoCosm (*37*) by performing BLASTP searches using as queries the FCP sequences from *C. gracilis* (*113*, *114*) and *Ectocarpus siliculosus* (*114*). Searches were run with BLOSUM62 and an E-value cutoff of 1.0×10⁻⁵ (relaxed to 1.0×10⁻³ where necessary to capture divergent candidates). In addition, transcriptomic data from *M. pyrifera* (*36*) were screened with the same queries and parameters, which revealed additional FCP sequences not annotated in the genome. Altogether, this procedure yielded 57 non-redundant FCP sequences across genome and transcriptome resources. For subsequent phylogenetic analysis, sequences that were incomplete, truncated or showed ambiguous annotation were removed.

We subjected the above set of M. pyrifera FCP genes to a phylogenetic analysis to determine the FCP sub-family composition and topology. To provide phylogenetic context and an outgroup for rooting, we added FCP/LHC sequences previously identified and validated from *E. siliculosus*, *C. gracilis* and *T. pseudonana* (*114*). To root the tree, we used a curated set of canonical green-algal LHCs. Specifically, four reference sequences were included: LHC from *Volvox africanus* (UniProt ID: A0A8J4EUW6), LHC from *Volvox reticuliferus* (UniProt ID: A0A8J4LST4), LHC from *Chlorella vulgaris* (UniProt ID: A0A9D4TGL7) and LHC from *Chlamydomonas reinhardtii* (UniProt ID: A8IKC8). This set was used exclusively for tree rooting to provide a stable outgroup reference. All sequences were aligned with MAFFT on the EBI web server using the L-INS-i strategy, BLOSUM62, a gap extension/offset penalty set to 0.123 and a maximum of 100 iterative refinement cycles. Misaligned or compositionally biased regions were corrected by manual inspection in Aliview v1.30 (*115*). The alignment file was used to reconstruct phylogenetic trees. Maximum-likelihood (ML) phylogenies were inferred with PhyML 3.0 (*116*) under the best-fit amino-acid substitution model selected by Smart Model Selection using AIC (*117*). Branch support was quantified with the SH-like approximate likelihood ratio test (SH-aLRT) and the approximate Bayes test (aBayes) as implemented in PhyML 3.0 (*116*). Trees were rooted on the green-algal LHC clade. Branch lengths are reported as substitutions per site. Resulting topologies were visualized and minimally edited to adjust tip label orientation and improve overall readability in FigTree v1.4.4 (*118*).

### FCP subfamily assignment

To assign subfamilies to the *M. pyrifera* FCPs with an assigned protein sequence (FCP1/3-11/13/17), subfamilies were determined based on the location of the sequence in *M. pyrifera*’s phylogenetic tree. For the *M. pyrifera* FCPs without an assigned sequence (second-belt FCP2/15/16/19/A/B, poly-alanine models), we used map-model correlation scores to identify the best-matching family. Models of each of the assigned families were fit into the focused-refined maps of each unassigned FCP, using the ChimeraX command fitmap, with manual adjustment as needed. The models used to represent the subfamilies were *M. pyrifera*’s FCP6 for Lhcr, FCP8 for Lhcq, FCP1 for RedCAP, FCP3 for CgLhcr9-like, FCP17 for Lhcf. To determine the quality of the fit, the Q-score between each model and each map was determined using the Q-score command in the ChimeraX toolshed. Using this method, we determined the best-fit subfamily of FCP15/16/19/A as Lhcf. Subfamilies for FCP2/B were not assigned.

## Acknowledgments

We thank Monterey Bay Seaweeds for providing kelp samples for our research, and S. Nuzhdin for providing access to the *M. pyrifera* genome prior to publication. We thank D. Barty, K. Bejar, V. Dubinin, Y. Xiang, H. Tiet, H. Wimboeck, C. Wang and M. Xu for kelp organelle isolations, C. Richards for lab management, BIOEM Director F. Guo and S2C2 staff members I. Fries and A. Cassago for technical support. We thank members of the Lagarias and Letts labs, especially M. Ayala-Hernandez, for advice, shared materials and equipment. We thank J. del Mármol, J. Letts, N. Rockwell, S. Theg and members of the Letts lab for comments on the manuscript. Some of this work was performed at the Stanford-SLAC Cryo-EM Center (S2C2), which is supported by the National Institute of General Medical Sciences (1R24GM154186). The content is solely the responsibility of the authors and does not necessarily represent the official views of the National Institutes of Health.

## Author contributions

Conceptualization: MM

Methodology: PM, JW, RR, GW, PZ, MM

Investigation: PM, JW, HMO, RR, GW, MM

Visualization: PM, JW, HMO, GW, VD, PZ, MM

Funding acquisition: MM

Project administration: MM

Supervision: PZ, MM

Writing – original draft: PM, JW, MM

Writing – review & editing: all authors

## Data and materials availability

The atomic coordinates for the *M. pyrifera* PSI-FCP supercomplex have been deposited (PDB: 9YGV). Maps corresponding to the composite map (EMDB: 72770) as well as 19 focused-refinement maps for each component of the composite map have been deposited. Details are available in **table S2**. The raw cryoEM data has been deposited (EMPIAR:12998).

## Supplementary Materials

### Supplementary text

#### Structural analysis of M. pyrifera*’s* PSI and predicted electron donors suggest a Type-III mechanism of electron transfer

PSI receives electrons from soluble electron donors that bind on the lumenal surface. These electrons reduce the photo-oxidized special pair P700 at the reaction center and are then transferred through cofactors to ferredoxin, which binds on the PSI stromal domain (**fig. S8A**). The electron donor interacts with the N-terminal α1 helix of PsaF *via* a long-range interactions before docking at the PsaA/B interface (*119*) (**fig. S8B-C**). Two families of electron donors have been described for PSI: plastocyanin (Pc), a copper-binding protein widely used in the green lineage, and cytochrome *c*_6_ (cyt *c*_6_), a *c*-type heme protein typically found in cyanobacteria and the red lineage (*120*, *121*). Pc appears to have been lost in most red algae and the red lineage (*121*). However, Pc has been seen to act as an alternative PSI donor under iron limitation in certain diatoms (*122*). The *M. pyrifera* genome encodes both Pc and cyt *c*_6_ (*123*), raising questions about the preferred electron donor and transfer mechanism to PSI. To examine how Pc and cyt *c*_6_ might interact with PSI, we identified their coding sequences in the *M. pyrifera* genome, generated structural-prediction models and docked the models onto *M. pyrifera*’s PSI.

As seen in sequence alignments and electrostatic-surface analysis, similar to plants, green algae and diatoms, the lumenal region of *M. pyrifera*’s PsaF possesses a positively charged insertion between α1 and α2 helices relative to cyanobacteria (**fig. S8D-F**). In *M. pyrifera*, both Pc (gene ID: DN119526 c0 g1 i1) and cyt *c*_6_ (gene ID: DN39228 c0 g1 i3) display conserved acidic patches that complement this positively charged region (*123*) (**fig. S8G-I**). Moreover, unbiased docking of predicted structural models of *M. pyrifera* Pc and cyt *c*_6_ to PSI positioned their cofactors within ∼15 Å (Pc) and ∼17 Å (cyt *c*_6_) of the reaction center (**fig. S8J-K**), with predicted binding energies of −5.7 kcal/mol (Pc) and −9.2 kcal/mol (cyt *c*_6_). These docking results resemble the hydrophobic docking interface to PsaA/PsaB subunits, the distance to the P700 special pair, and the orientation of the negatively charged donor surfaces toward the α1 helix of PsaF, as described in high-resolution structures of PSI-donor complexes from cyanobacteria and plants, as well as in diatom *in silico* models (*124*, *125*). The stronger interaction of *M. pyrifera*’s cyt *c*_6_ relative to Pc is consistent with the cyt *c*_6_ preference reported in diatoms, suggesting that brown algal PSI predominantly uses cyt *c*₆ while maintaining the capacity to interact with Pc (*125*). Moreover, the electrostatic complementarity between the basic patch on PsaF and the acidic surfaces of Pc and cyt *c*_6_ suggests that, like most eukaryotic phototrophs, *M. pyrifera* uses a long-range, electrostatics-based Type-III mechanism for electron transfer to PSI, rather than Type I or II collision-based mechanism described in cyanobacteria (*125–127*).

#### Stromal domain: implications for ferredoxin (Fd) binding

PSI transfers electrons to ferredoxin (Fd), which then acts as a central reducing agent in chloroplasts, including the reduction of NADP^+^ to NADPH by FNR (*1*). The PSI:Fd electron transfer is enabled by PsaC’s 4Fe4S clusters (**fig. S8A**). In previously studied organisms, the protein:protein interactions between PSI and Fd are mediated by electrostatic interactions with PsaA/C/D/E (*35*). Upon examination of the *M. pyrifera* stromal domain, we determined that all reported residues mediating the PSI:Fd interaction are conserved, except for a substitution of Arg to Lys in *Mp*PsaE-K4 and the loss of Lys to Thr in *Mp*PsaD-T105, both of which are conserved in other brown algae (*35*) (**fig. S8L**).

The recent diatom PSI structures revealed the presence of an additional subunit (PsaS, a.k.a. Psa29) in the stromal domain (*28*, *30*). Diatom PsaS interacts mainly with PsaC and PsaD, with some additional minor interactions with stroma-facing regions of PsaB/I/L. Given that PsaS does not contain redox cofactors and that its binding site is ∼14 Å away from Fd at its closest, it is unlikely that PsaS has direct effects on Fd recruitment or electron transfer. Rather, PsaS may serve to stabilize PSI’s stromal domain or to indirectly regulate the PSI:Fd interaction. In contrast to diatoms, we did not find evidence for a PsaS homologue in the *M. pyrifera* cryoEM density, even after specific masked classifications and focused refinements of this area using a diatom PsaS models (*28*, *30*). We also failed to find homologues PsaS using diatom PsaS queries in homologue searches in *M. pyrifera*, *E. siliculosus* or Phaecyacea at large. This is in line with the finding (*30*) that PsaS is absent in Bolidophyceae, a sister group to diatoms. Further, it suggests that PsaS is a diatom innovation not present in the common ancestor of Ochrophyta. By extension, this implies that Chrysista and Diatomista clades use different mechanisms to regulate electron transfer from PSI to Fd. Evaluation of electron transfer in the presence and absence of PsaS is needed to test this hypothesis.

### Supplementary methods

#### Identification of plastocyanin and cytochrome c₆ coding sequences

Coding sequences for Pc and cyt *c*₆ were identified in the *Macrocystis pyrifera* genome using tBLASTn searches against the PhycoCosm database (Macrocystis pyrifera CI_03 v1.0). For Pc, the *Synechocystis sp.* PCC 6803 sequence (UniProt: P21697) was used as query, while for cyt *c*₆ the *Ectocarpus siliculosus* sequence (UniProt: A0A6G6D7E1) was employed. Searches were perfomed using with an expectation E-value 1.05 × 10⁻⁵ and the BLOSUM62 substitution matrix. We used the *M. pyrifera* “all-model-transcripts” dataset (release 202220914). Candidate coding sequences were translated into protein sequences using SnapGene to determine the corresponding open reading frames and confirm the presence of full-length proteins. Signal peptides were predicted with SignalP 6.0 (*128*), and mature protein sequences were extracted for downstream analysis.

#### Sequence alignment and electrostatic analysis of PsaF, cyt c_6_, and Pc

PsaF sequences included *Synechocystis sp.* PCC 6803 (UniProt: P29256), *C. gracilis* (UniProt: A0A345U7L1), *G. sulphuraria* (UniProt: E3UIU5), and *Pisum sativum* (UniProt: E3UIU5). Cytochrome *c*₆ sequences were aligned from *M. pyrifera* (predicted, gene ID: DN39228 c0 g1 i3), *Phaeodactylum tricornutum* (UniProt: B5Y578) and *Synechocystis sp.* PCC 6803 (UniProt: P46445). Plastocyanin alignments included *M. pyrifera* (predicted, gene ID: DN119526 c0 g1 i1), *P. sativum* (UniProt: P16002), and *Synechocystis sp.* PCC 6803 (UniProt: P21697). All multiple sequence alignments were performed using ClustalW with standard parameters.

#### Structural modeling of electron donors and molecular docking

Structural prediction models of Pc and cyt *c*₆ were generated with AlphaFold3, incorporating the copper ion in Pc and the heme C group in cyt *c*₆. The top-ranking models were selected based on pLDDT confidence scores and stereochemical quality metrics from MolProbity. Conserved metal-binding motifs were confirmed by comparison to homologous protein structures (PDB: 1AG6 for Pc and 1CYI for cyt *c*_6_). Electrostatic potential surfaces of PsaF, Pc and cyt *c*_6_ were obtained using AMBER20 in ChimeraX. Docking simulations were performed between the *M. pyrifera* PSI cryo-EM structure and AlphaFold-predicted Pc and cyt *c*₆ models using HADDOCK 2.4 (*129*). The protocol included rigid-body docking, semi-flexible refinement. The best-scoring clusters were selected according to HADDOCK scores and interface consistency with known PSI-donor complexes from cyanobacteria and plants. Final docking poses were analyzed in ChimeraX to measure donor–P700 distances. Binding free energies were estimated using the PISA server (*130*).

**Fig. S1.**
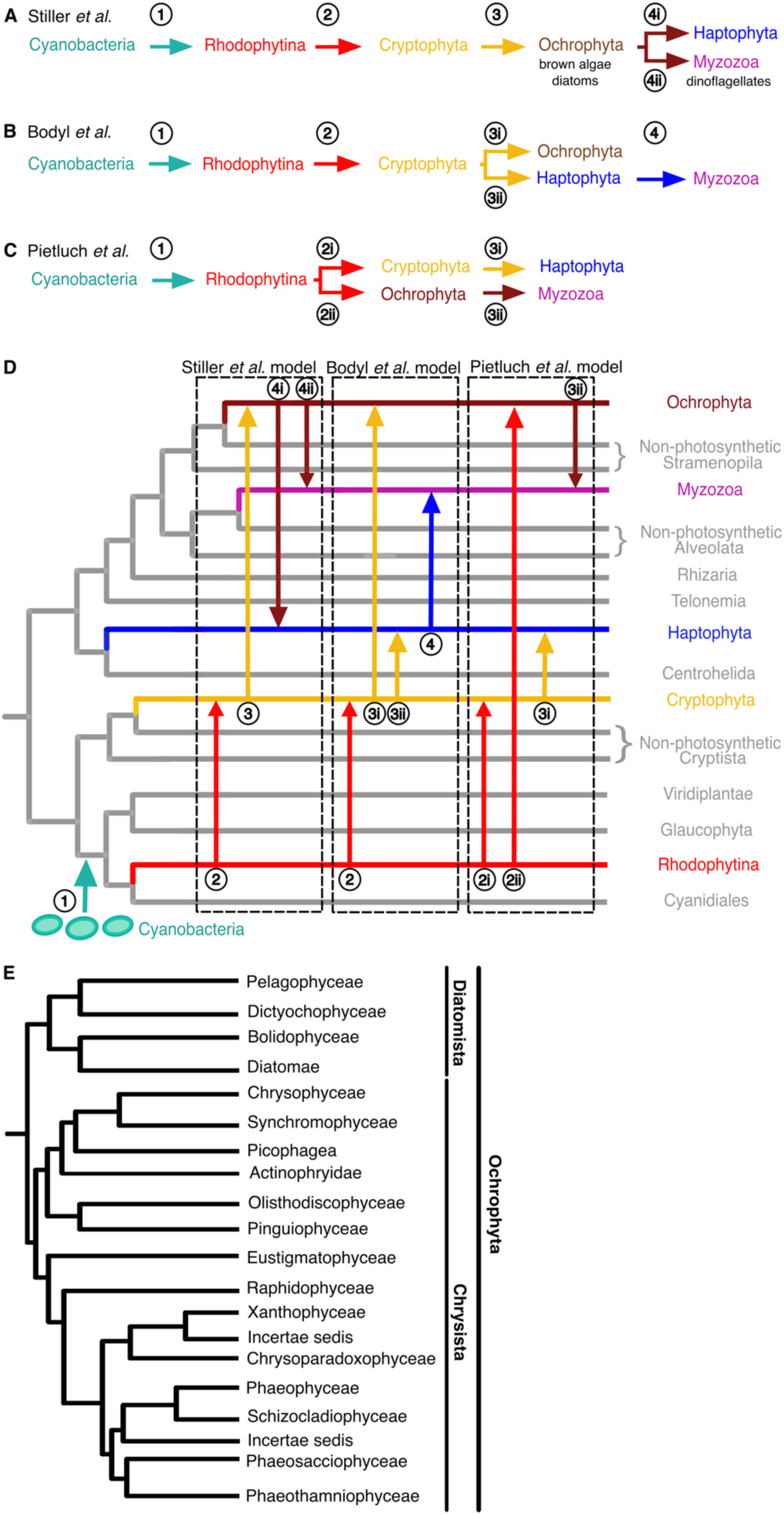
Evolutionary relationships in the red lineage. (**A-C**) Models of endosymbioses in the red lineage consistent with molecular timescale analysis (*14*, *15*) proposed by Stiller *et al*. (*85*) (A), Bodyl *et al*. (*86*) (B) and Pietluch *et al.* (*15*)(C). (D) Representation of (A-C) over cladogram. Arrows and numbers represent serial and parallel endosymbioses. (E) Cladogram of Ochrophyta phylum (*62*).

**Fig. S2.**
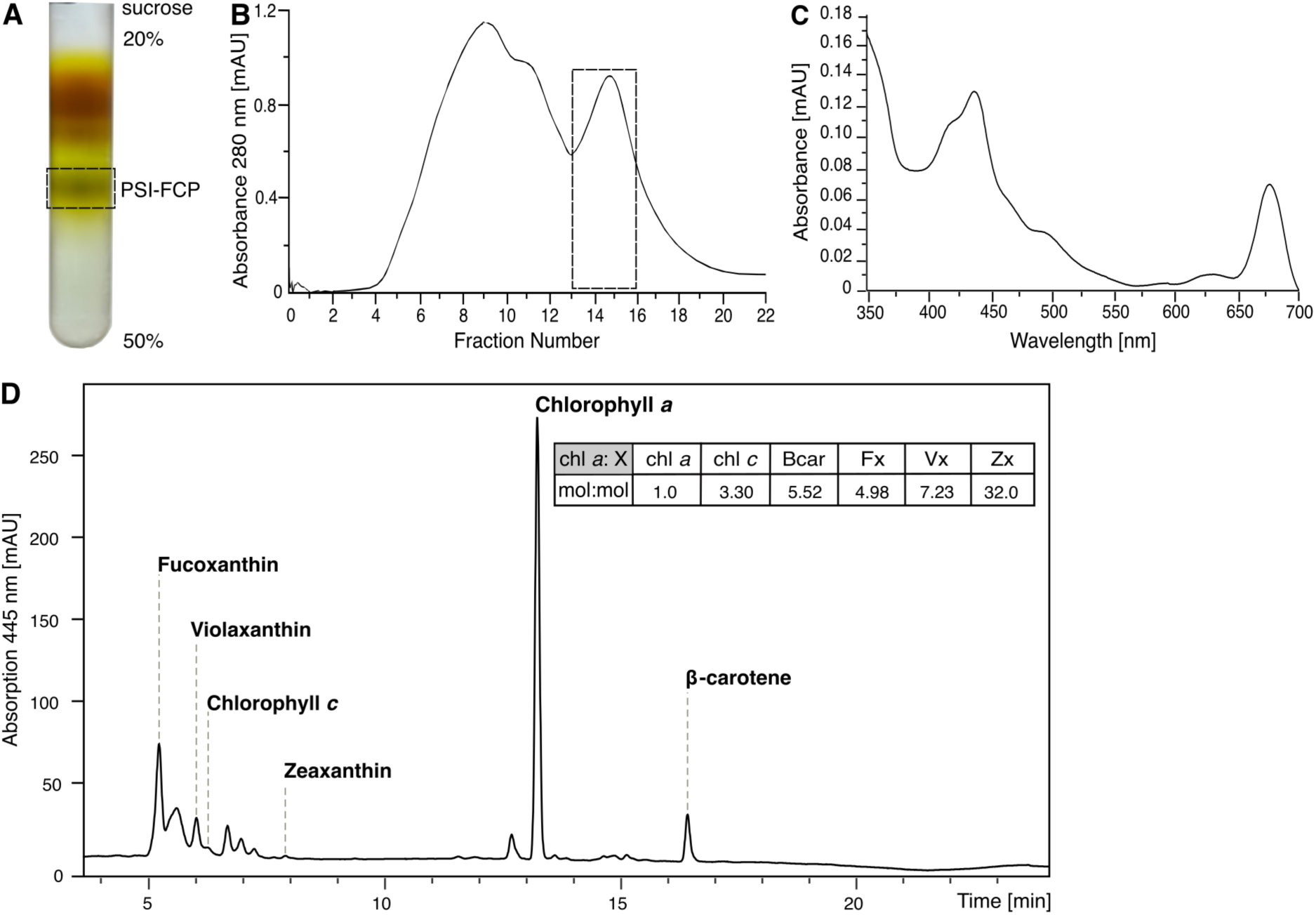
Purification and biochemical characterization of *Macrocystis pyrifera*’s PSI-FCP supercomplex. (**A**) Linear sucrose gradient (20-50%, w:v) before fractionation. (**B**) Chromatogram of the fractions from (A), showing the absorbance monitored at 280 nm. Fractions that were selected and pooled for subsequent cryoEM processing are indicated in a dashed box. (**C**) Absorption spectrum monitored at 350-750 nm of the pooled purified fractions indicated in (B). (**D)** Chromophore analysis of purified fractions using high-performance liquid chromatography (HPLC) based on absorption at 445 nm. Chromophores were identified by comparison to authentic standards. Inset table shows ratios (mole:mole) of chl *a* relative to the other chromophores. chl *a*, chlorophyll *a*; chl *c*, chlorophyll *c*; BCar, ß-carotene; Fx, fucoxanthin; Vx, violaxanthin; Zx, zeaxanthin.

**Fig. S3.**
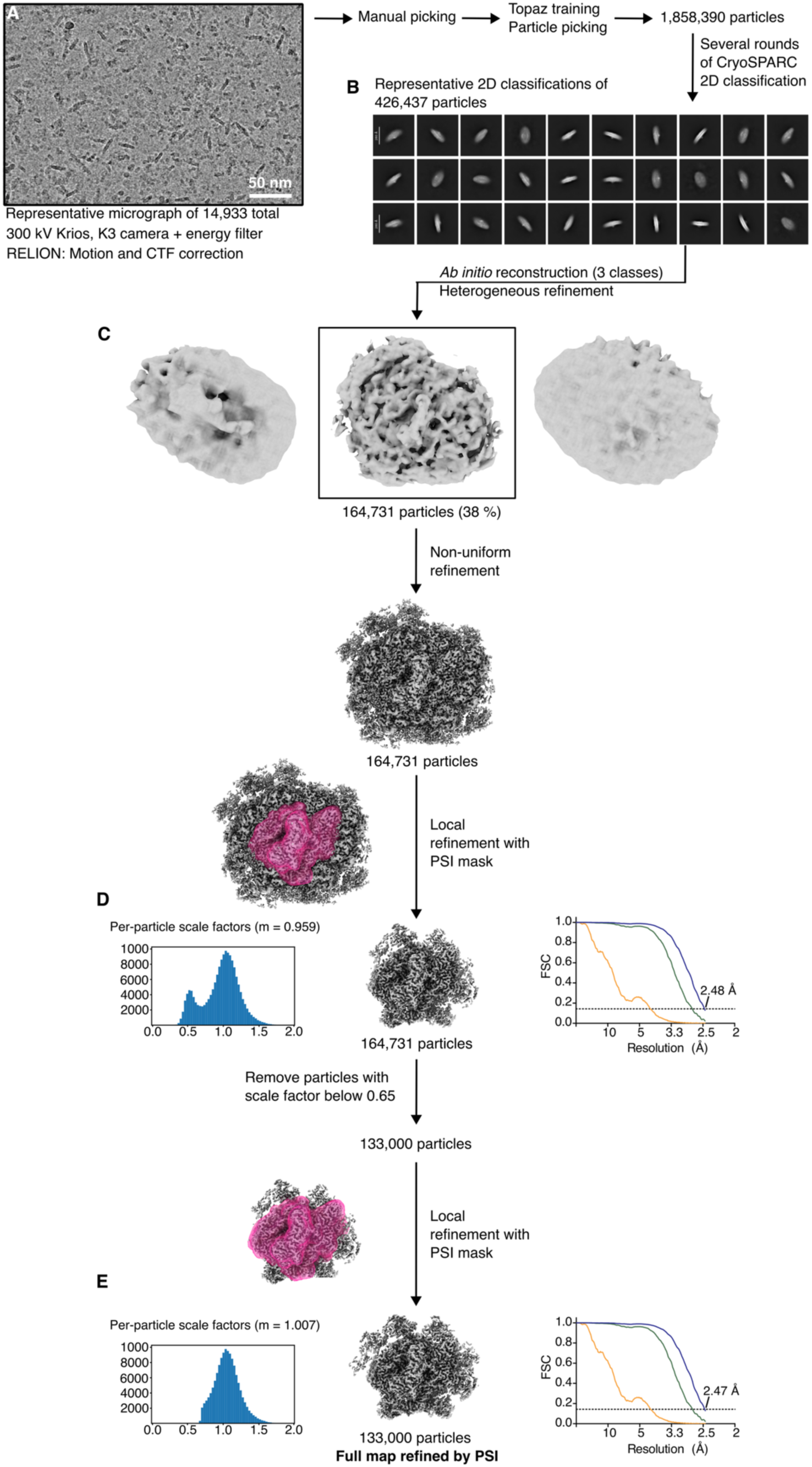
CryoEM micrograph and initial data processing for *Macrocystis pyrifera*’s PSI-FCP supercomplex. (**A**) Representative micrograph of 14,933 collected. (**B**) Representative 2D classes of PSI-FCP particles. (**C**-**E**) Initial 3D classification and refinement pipeline to obtain the full map refined by PSI (E). The Fourier shell correlation (FSC) curves are shown for no (orange), loose (green) and tight (blue) masking. The resolution at which the tight masked FSC crosses the 0.143 gold-standard limit (dashed line) is indicated. Per-particle scale factor distribution for the intermediate (D) and final (E) maps are also shown. Note that the final map refined by PSI (E) was used as the starting point for the focused refinement of the individual FCPs shown in **Fig. S4-6**.

**Fig. S4.**
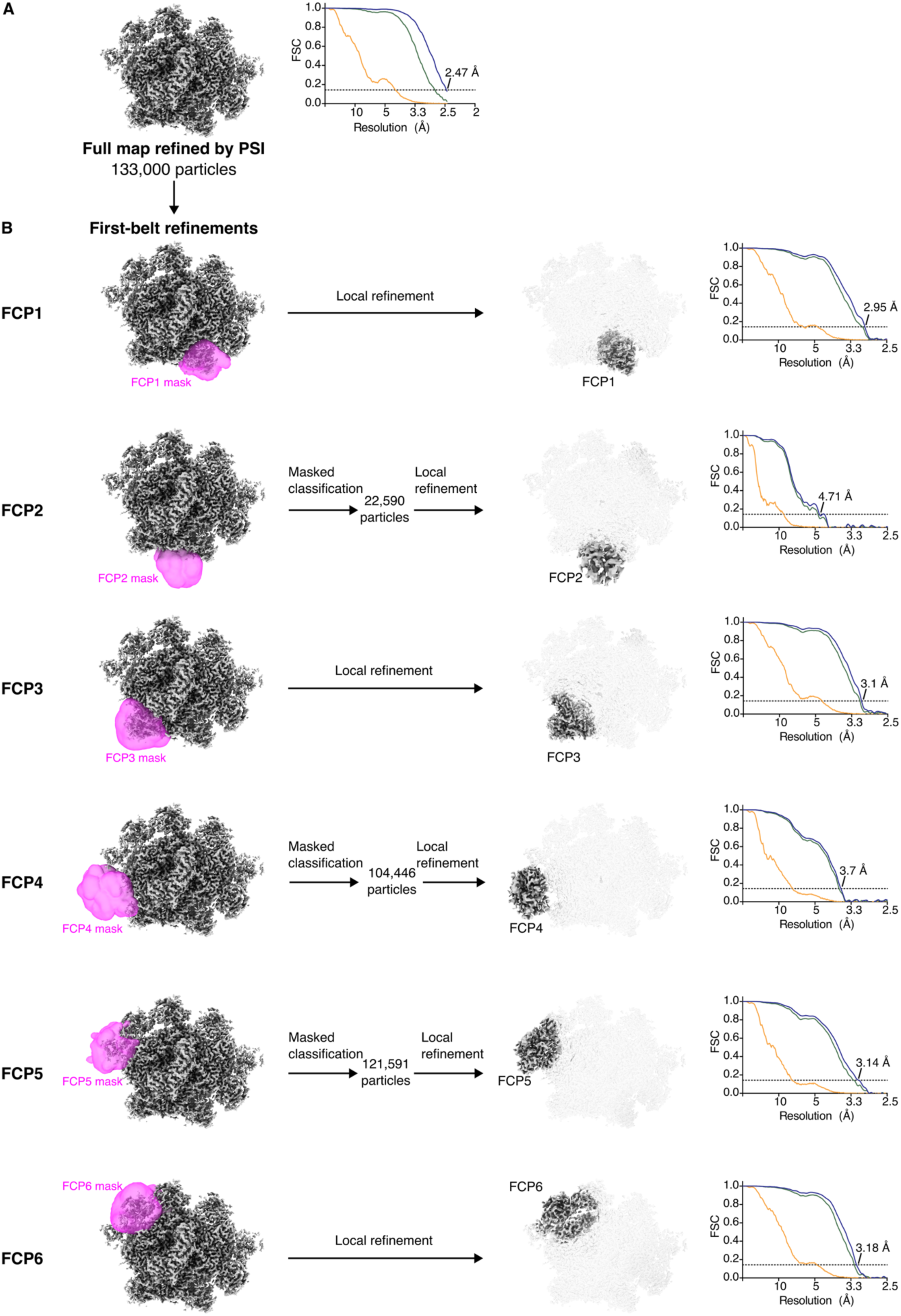
Masked classification and focused refinements of FCP1-FCP6 (first belt). (**A**) The starting map for the focused refinements was the map refined by PSI and its corresponding particle stack, shown in **Fig. S3**. (**B**) Focused refinements using individual masks (magenta) were performed to improve the map quality of FCP1-6. Due to sub-stoichiometric occupancy of the FCP2/4/5/6, a masked classification using the corresponding individual mask was performed before the focused refinement of those specific regions, using the same mask. Fourier shell correlation (FSC) curves for the individual focused refinements are shown for no (orange), loose (green) and tight (blue) masking. The resolution at which the tight mask FSC crosses the 0.143 gold-standard limit (dashed line) is indicated.

**Fig. S5.**
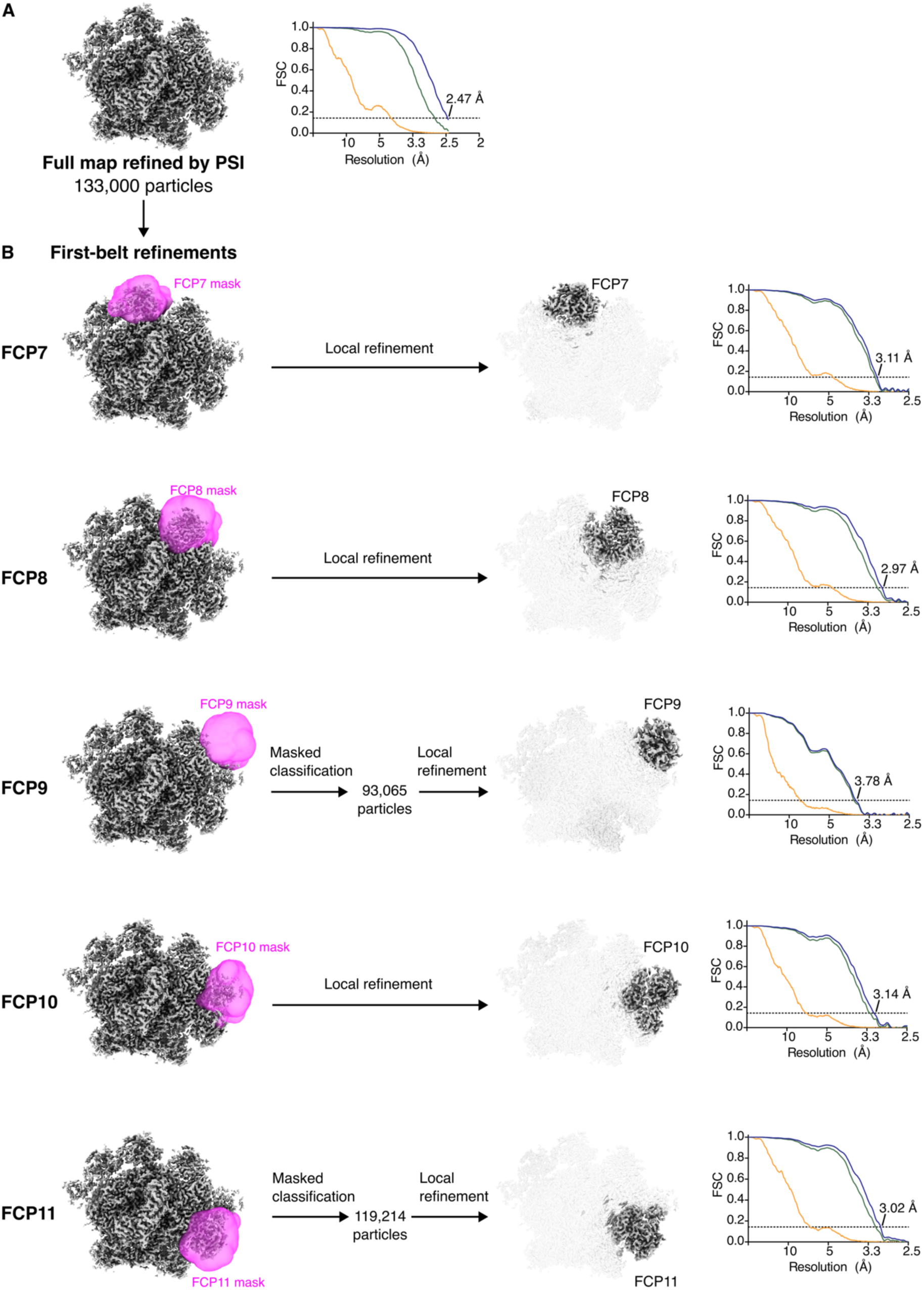
Masked classification and focused refinements of FCP7-FCP11 (first belt). (**A**) The starting map for the focused refinements was the map refined by PSI and its corresponding particle stack, shown in **Fig. S3**. (**B**) Focused refinements using individual masks (magenta) were performed to improve the map quality of FCP7-11. For FCP9/11, a masked classification using the corresponding individual mask was performed before the focused refinement of those specific regions, using the same mask. Fourier shell correlation (FSC) curves for the individual focused refinements are shown for no (orange), loose (green) and tight (blue) masking. The resolution at which the tight mask FSC crosses the 0.143 gold-standard limit (dashed line) is indicated.

**Fig. S6.**
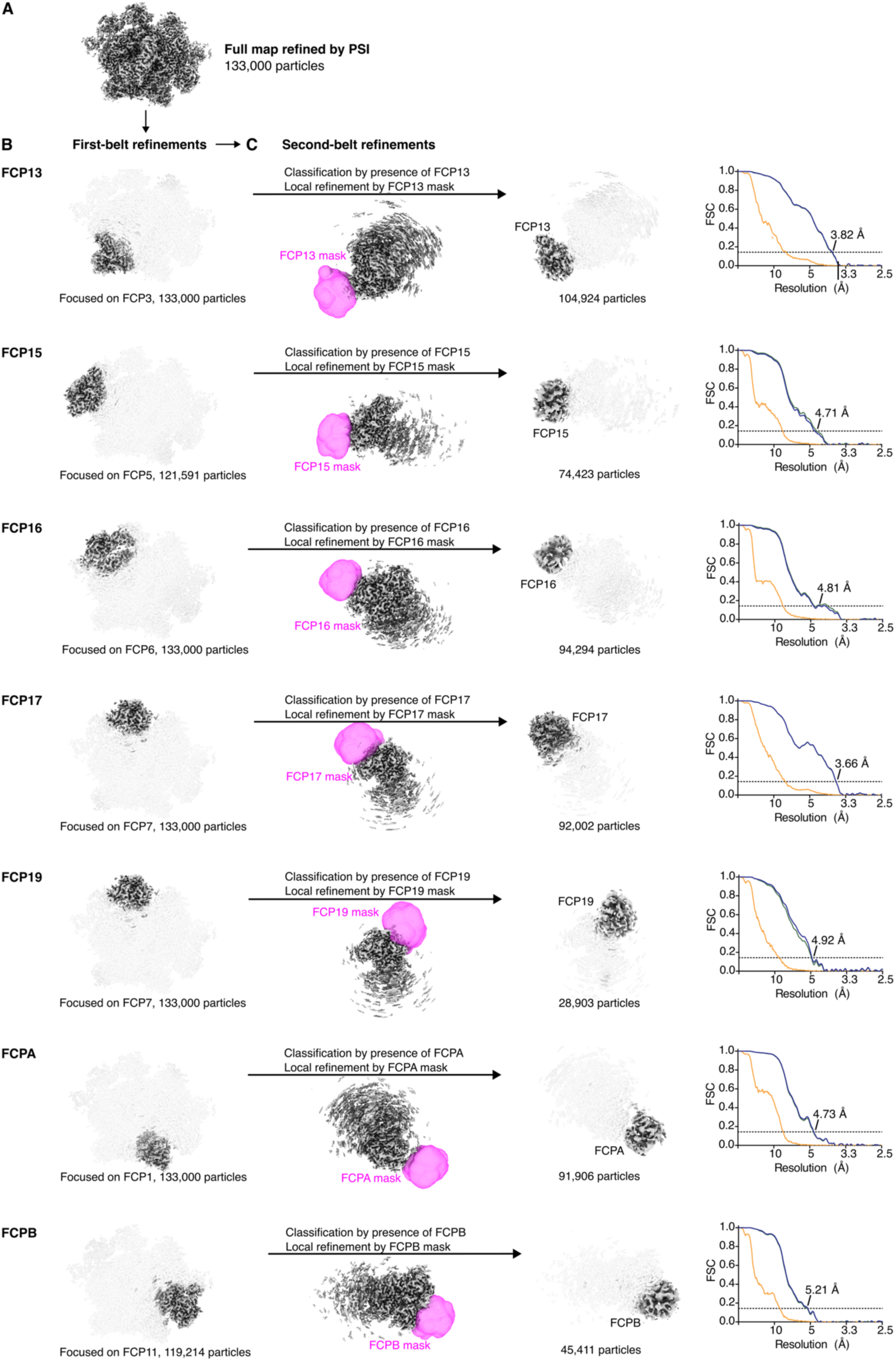
Masked classification and focused refinements of FCP13/15/16/17/19/A/B (second-belt). (**A**-**C)** Focused refinements were performed in three subsequent steps: (A) refinement by PSI, as shown in **Fig. S3**, (B) refinement by a relevant first-belt FCP, as shown in Fig. S3 and S4, (C) masked classification and refinement by the second-belt FCP, using the masks shown (magenta). Fourier shell correlation (FSC) curves for the individual focused refinements are shown for no (orange), loose (green) and tight (blue) masking. The resolution at which the tight mask FSC crosses the 0.143 gold-standard limit (dashed line) is indicated.

**Fig. S7.**
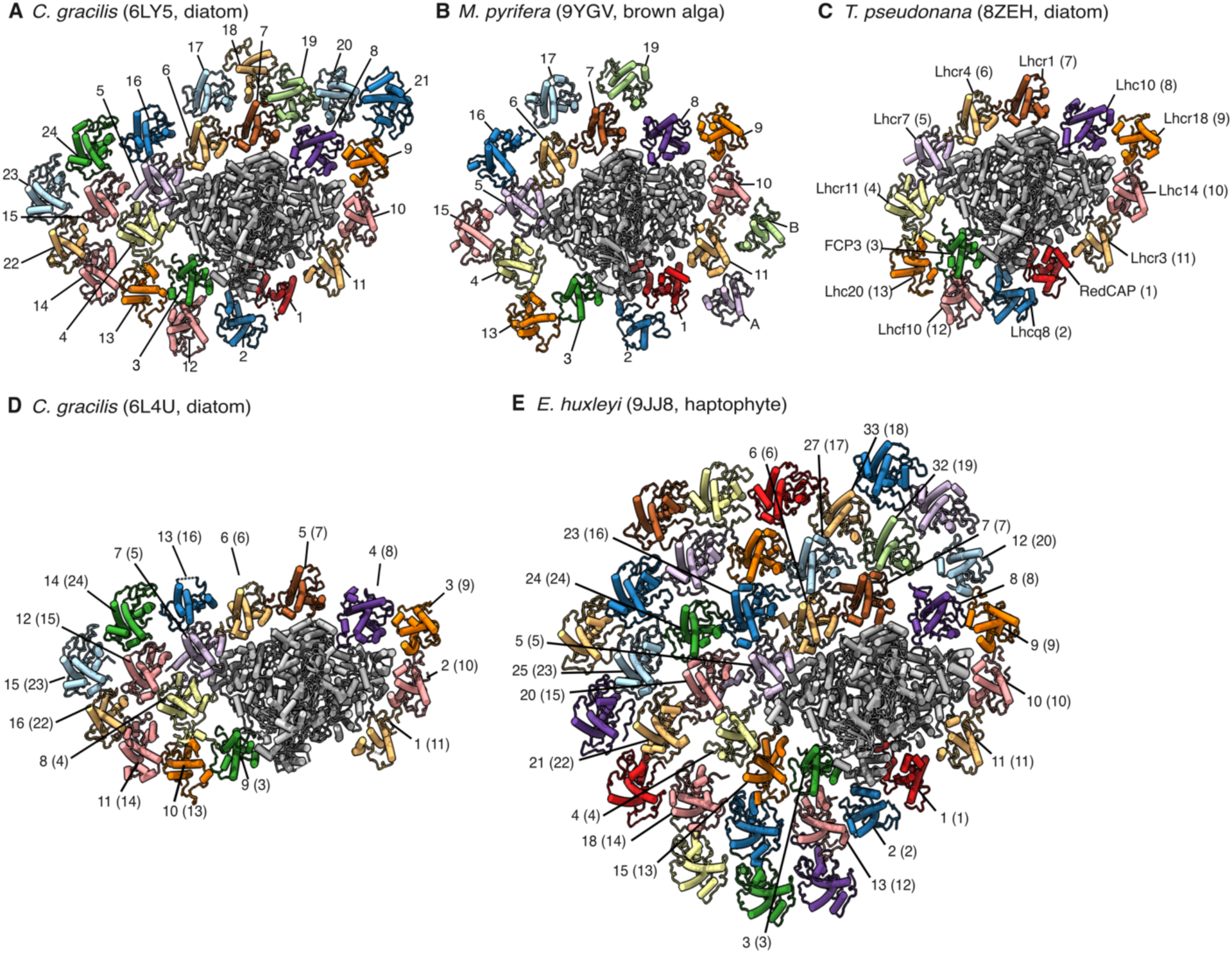
FCP antenna numbering and conversion. (**A-E**) PSI-FCP supercomplexes from (A) *Chaetoceros gracilis* (PDB: 6LY5) (*28*), (B) *Macrocystis pyrifera* (PDB:9YGV), this work, (C) *Thalassiosira pseudonana* (PDB: 8ZEH) (*31*), (D) *C. gracilis* (PDB: 6L4U) (*29*), (E) *Emiliania huxleyi* (PDB: 9JJ8) (*25*). PDB codes shown next to species name. PSI shown in grey cartoon. FCP antenna shown in cartoon colored by subunit. Labels show nomenclature used in respective papers, with the number used in this work in parentheses. FCP suffixes omitted for clarity.

**Fig. S8.**
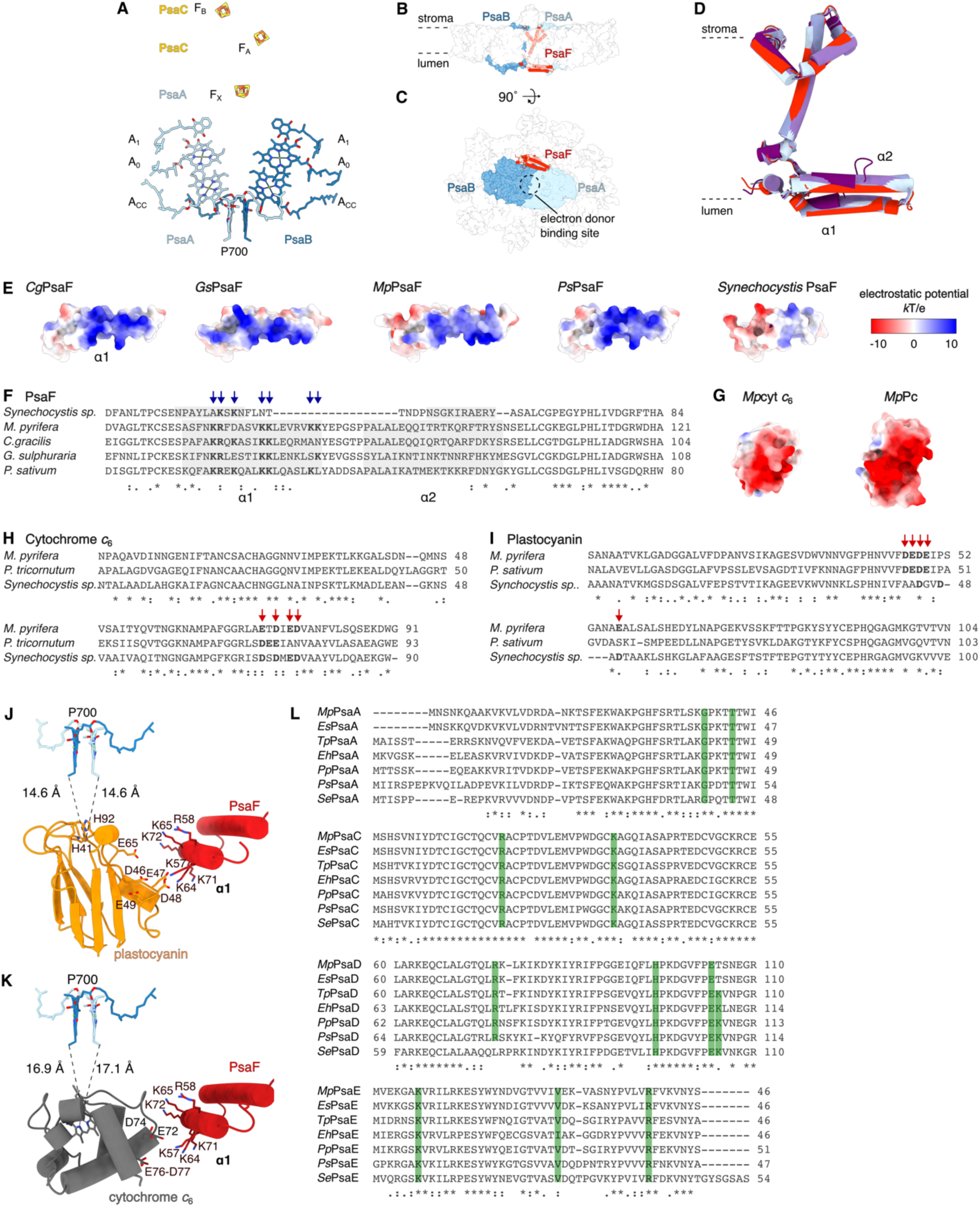
Predicted electron donor and acceptor regions of *M. pyrifera* PSI. (**A**) Detail of electron-transfer-chain co-factors in PsaA and PsaB shown in atom stick representation, colored by subunit. P700, chl special pair; A_CC_, accessory chl; A_0_, primary electron acceptor; A_1_, secondary electron acceptor; F_X_, F_A_, and F_B_, iron-sulfur clusters. (**B-C**) Membrane (B) and lumenal (C) views of *M. pyrifera* PSI-FCP in surface representation. PsaA/B/F colored by subunit; PsaA/B in surface; PsaF in cartoon. Approximate location of electron-donor binding site marked in dashed circle. (**D-F**) PsaF homolog comparisons. (D) Structural superposition *Chaetoceros gracilis* (light blue, PDB: 6L4U), *Galdieria sulphuraria* (red alga, dark blue, PDB:9KC5), *M. pyrifera* (red, PDB: 9YGV), *Pisum sativum* (light violet, PDB: 4XK8), and *Synechocystis sp.* (dark violet, PDB:5OY0). (**E**) Electrostatic surface of PsaF-α1 helix from *G. gracilis*, *G. sulphuraria gracilis*, *M. pyrifera*, *P. sativum*, and *Synechocystis sp*. Red, negative; white, neutral; blue, positive. (**F**) Sequence alignment of the N-terminal region of PsaF from *Synechocystis sp*., *M. pyrifera*, *C. gracilis*. *G. sulphuraria*, *P. sativum*, with α1–α2 helices (grey boxes) and conserved basic residues (blue arrows). (**G**) Electrostatic surfaces of predicted models of *M. pyrifera* (Mp) cytochrome *c*_6_ (cyt *c*_6_) and plastocyanin (Pc). Scale as in (E). (**H-I**) Sequence alignments of cyt *c*_6_ from *M. pyrifera*, *P. tricornutum*, and *Synechocystis sp*. PCC 680 (H) and Pc from *M. pyrifera*, *P.sativum*, and *Synechocystis sp*. PCC 6803 (I). Conserved acidic residues at docking interface marked with red arrows. (**J-K**) Docking models of *M. pyrifera* Pc (J) and cyt *c*_6_ (K) bound to *M. pyrifera* PSI. Distances from the redox centers of Pc and cyt *c*_6_ to special pair P700 are shown. (**L**) Sequence alignments of regions of PsaA, PsaC, PsaD and PsaE, implicated in ferredoxin binding from *M. pyrifera* (brown alga, *Mp*), *Ectocarpus siliculosus* (brown alga, *Es*), *Thalassiosira pseudonana* (diatom, *Tp*), *Emiliania huxleyi* (haptophyte, *Eh*), *Porphyridium purpurea* (red alga, *Pp*), *P. sativum* (plant, *Ps*), and *S. elongatus* (cyanobacterium, *Se*). Conserved residues associated with ferredoxin binding marked in green boxes. Symbols underneath aligned residues: * fully conserved, : conservation between group of strongly similar properties,. conservation between group of weakly similar properties.

**Fig. S9.**
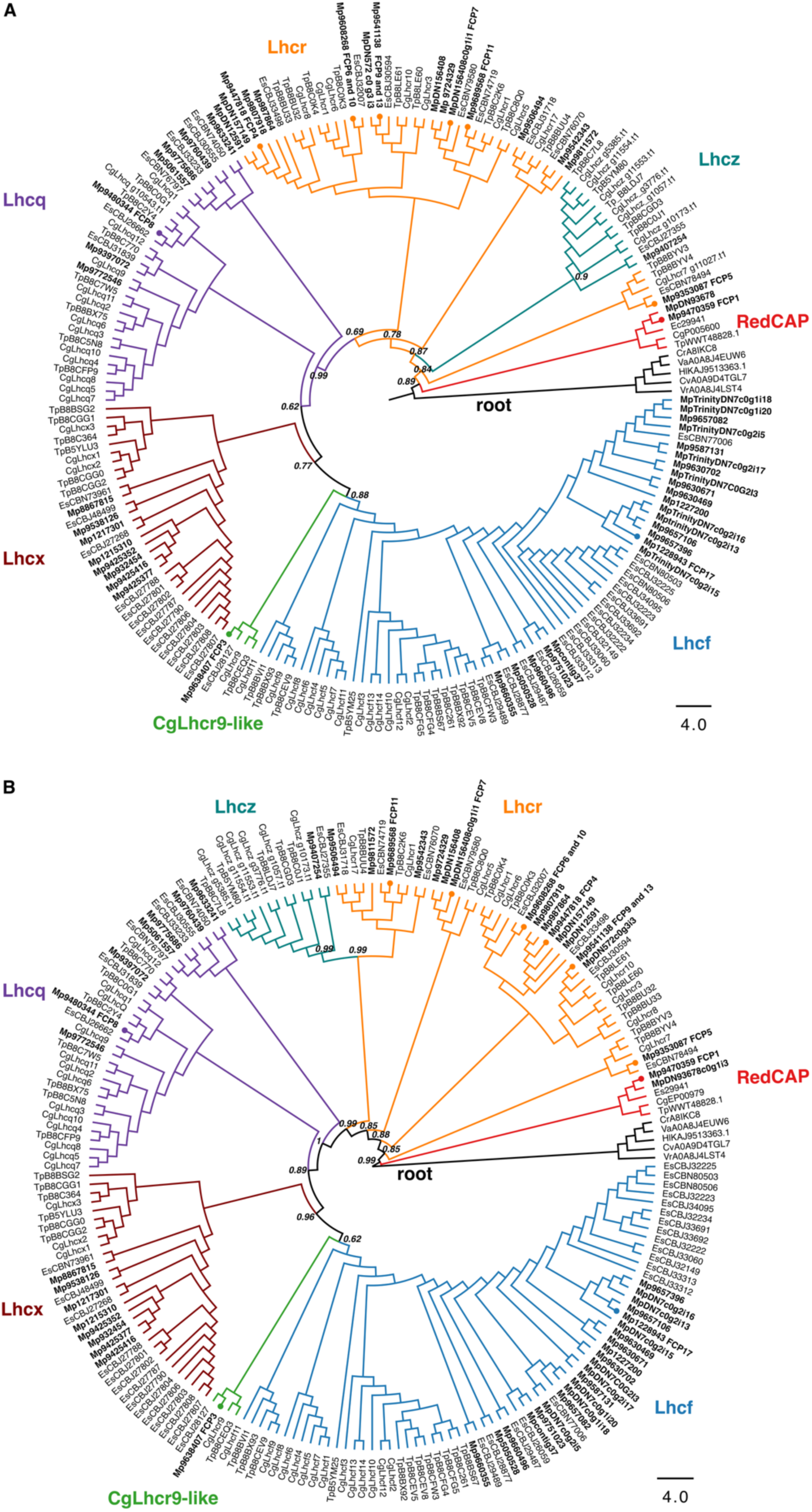
Phylogenetic analysis of ochrophyte FCP proteins. (**A-B**) Maximum likelihood phylogenetic tree with branch support values calculated using SH-aLRT test (A) or aBayes (B). Both trees were inferred from 203 LHC protein sequences from *Macrocystis pyrifera* (Mp), *Ectocarpus siliculosus* (Es), *Chaetoceros gracilis* (Cg), and *Thalassiosira pseudonana* (Tp). FCP subfamilies indicated by color: Lhcr (orange), Lhcf (blue), Lhcq (purple), Lhcx (brown), Lhcz (teal) and CgLhcr9-like (green); RedCAP of LHC superfamily in red. Black branches correspond to green algal LHC sequences used to root the trees. *M. pyrifera* sequences are highlighted in bold; those identified in the *M. pyrifera* PSI-FCP structure (PDB: 9YGV, this work) indicated with circle at branch terminus. Posterior probabilities supporting basal nodes that define each subfamily shown beside the node. Scale indicates number of substitutions per site. Sequence details are provided in **Supplementary Data File 1**.

**Fig. S10.**
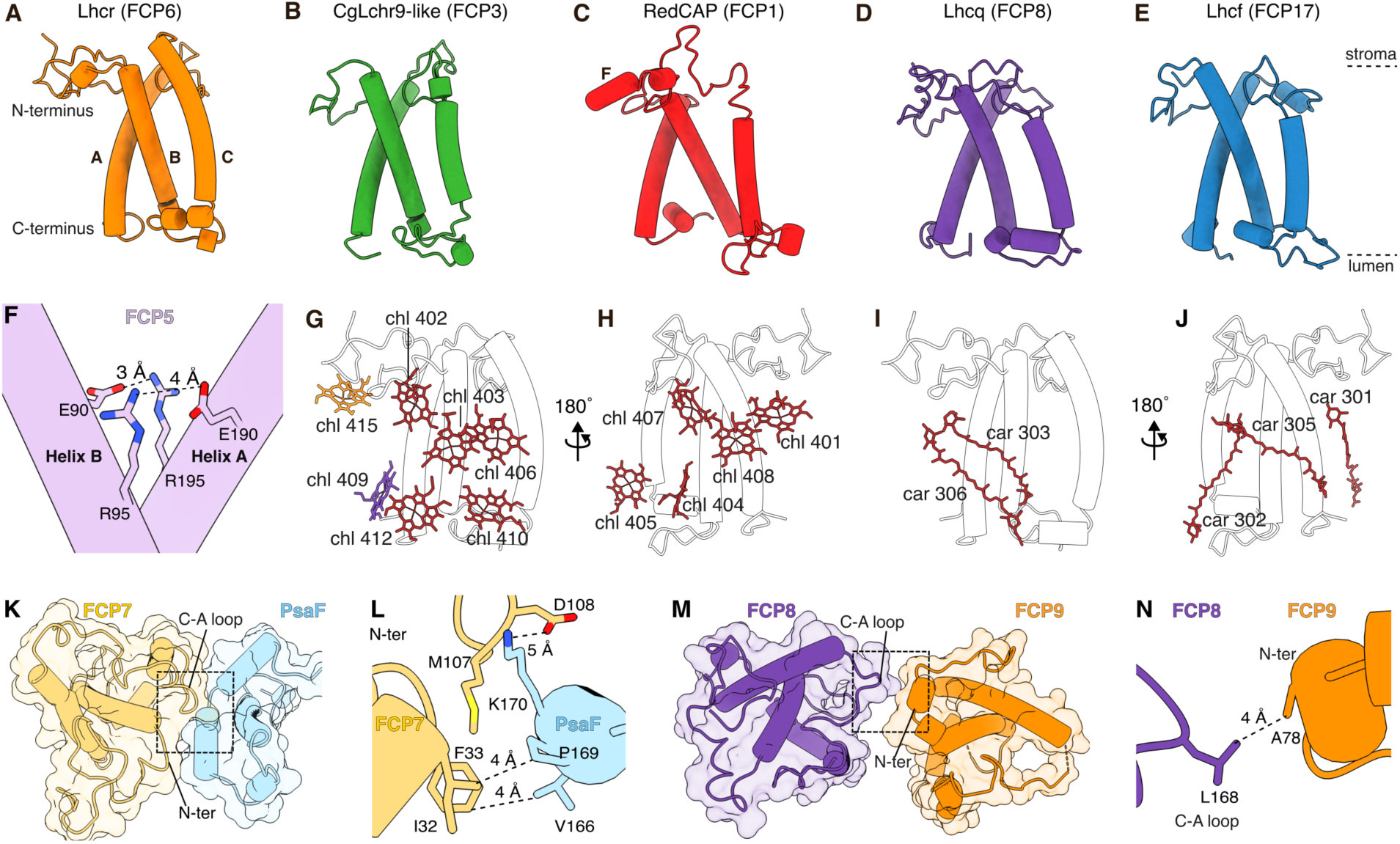
Basic structural characterization of *M. pyrifera*’s FCP subfamilies. (**A-E**) Representative structures of each subfamily shown in cartoon representation (name of actual FCP shown in parentheses), colored by subfamily. Helices A, B, C, as well as N- and C-termini labeled in (A). RedCAP’s helix F labeled in (C). Approximate positions of stroma and lumen marked with dashed lines. (**F**) Representative example (FCP5) of FCP Arg-Glu salt bridges between the A and B helices across subfamilies. (**G**-**J**) Full complement of the *M. pyrifera* FCP-chromophore binding sites shown for chlorophyll (G, H) and carotenoids (I, J) shown from the core-facing orientation (G, I) or outer-belt-facing orientation (H, J). Family-specific chromophores shown in the subfamily color (Lhcr in orange, RedCAP in red), non-family-specific shown in brown. (**K-L**) Representative example of first-belt FCP (FCP7) interacting with PSI subunit (PsaF) through the FCP C-A and N-terminal (N-ter) loops, viewed from the stroma. Subunits shown in cartoon with semi-transparent surface overlay, colored by subunit. Area in dashed box in (K) is detailed in (L) in slightly rotated angle. (**M, N**) Representative example of two first-belt FCPs interacting (FCP8/9) through the FCP A-C and N-terminal loops, viewed from the stroma. Subunits shown in cartoon with semi-transparent surface overlay, colored by subunit. Area in dashed box in (M) detailed in (N).

**Fig. S11.**
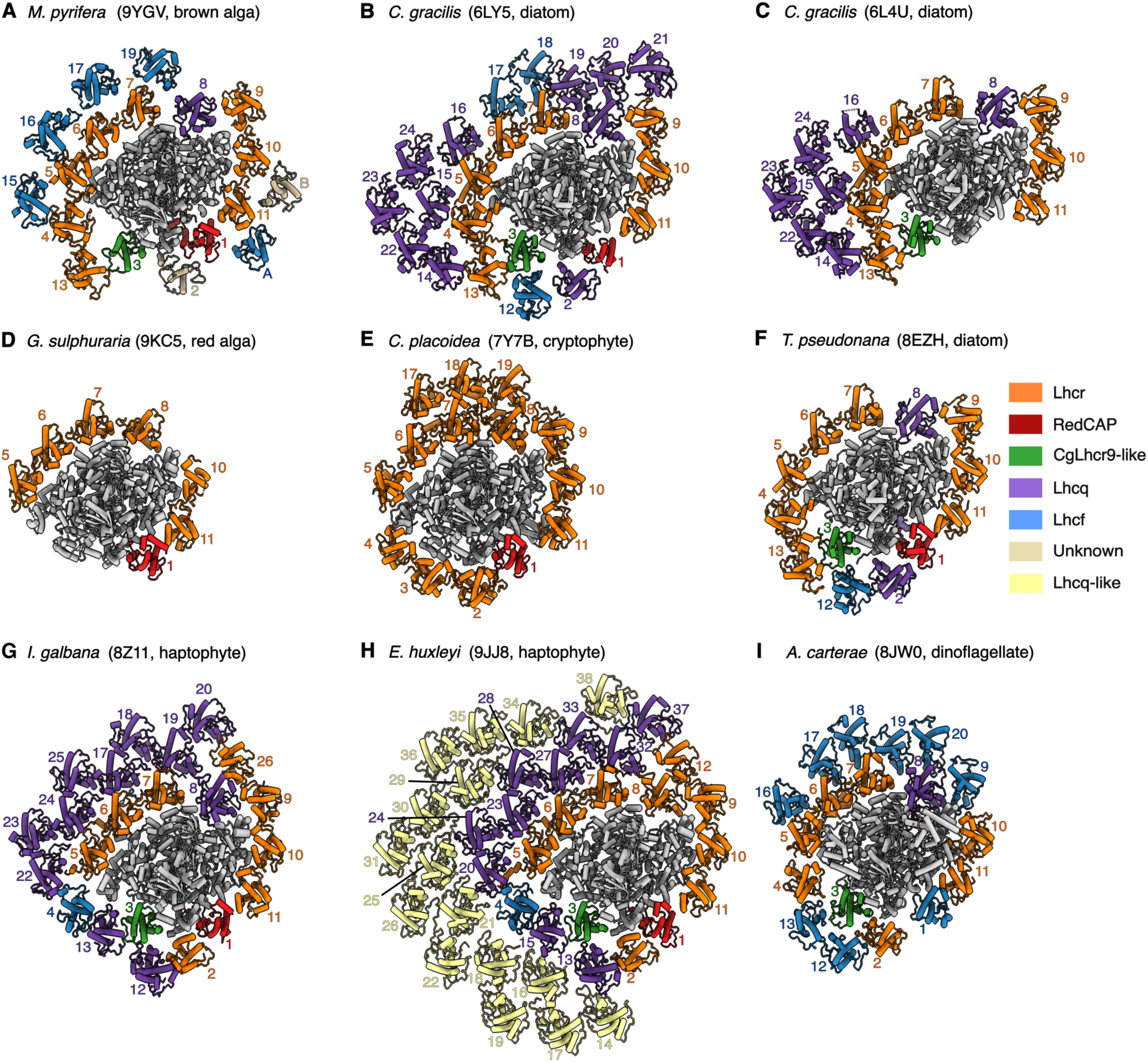
Comparison of FCP subfamilies in PSI antennae across red-lineage organisms. PSI-FCP supercomplexes from (**A**) brown alga *M. pyrifera* (PDB: 9YGV, this work), (**B**) diatom *C. gracilis* (PDB: 6LY5) (*28*), (**C**) diatom *C. gracilis* (PDB: 6L4U) (*29*), (**D**) red alga *G. sulphuraria* (PDB: 9KC5) (*18*), (**E**) cryptophyte *Croomonas placoidea* (PDB: 7Y7B) (*22*), (**F**) diatom *T. pseudonana* (PDB: 8ZEH) (*31*), (**G**) haptophyte *Isochrysis galbana* (PDB: 8Z11) (*24*), (**H**) haptophyte *Emiliania huxleyi* (PDB: 9JJ8) (*25*) and (**I**) dinoflagellate *Amphidinium carterae* (PDB: 8JW0) (*26*). PDB codes shown next to species name. PSI shown in grey cartoon. FCP antenna shown in cartoon colored by subunit, according to FCP subfamily: Lhcr in orange, RedCAP in red, CgLhcr9-like in green, Lhcq in purple, Lhcf in blue, unknown in beige, Lhcq-like in yellow. FCP numbers used in this work shown, with FCP suffixes omitted for clarity.

**Fig. S12.**
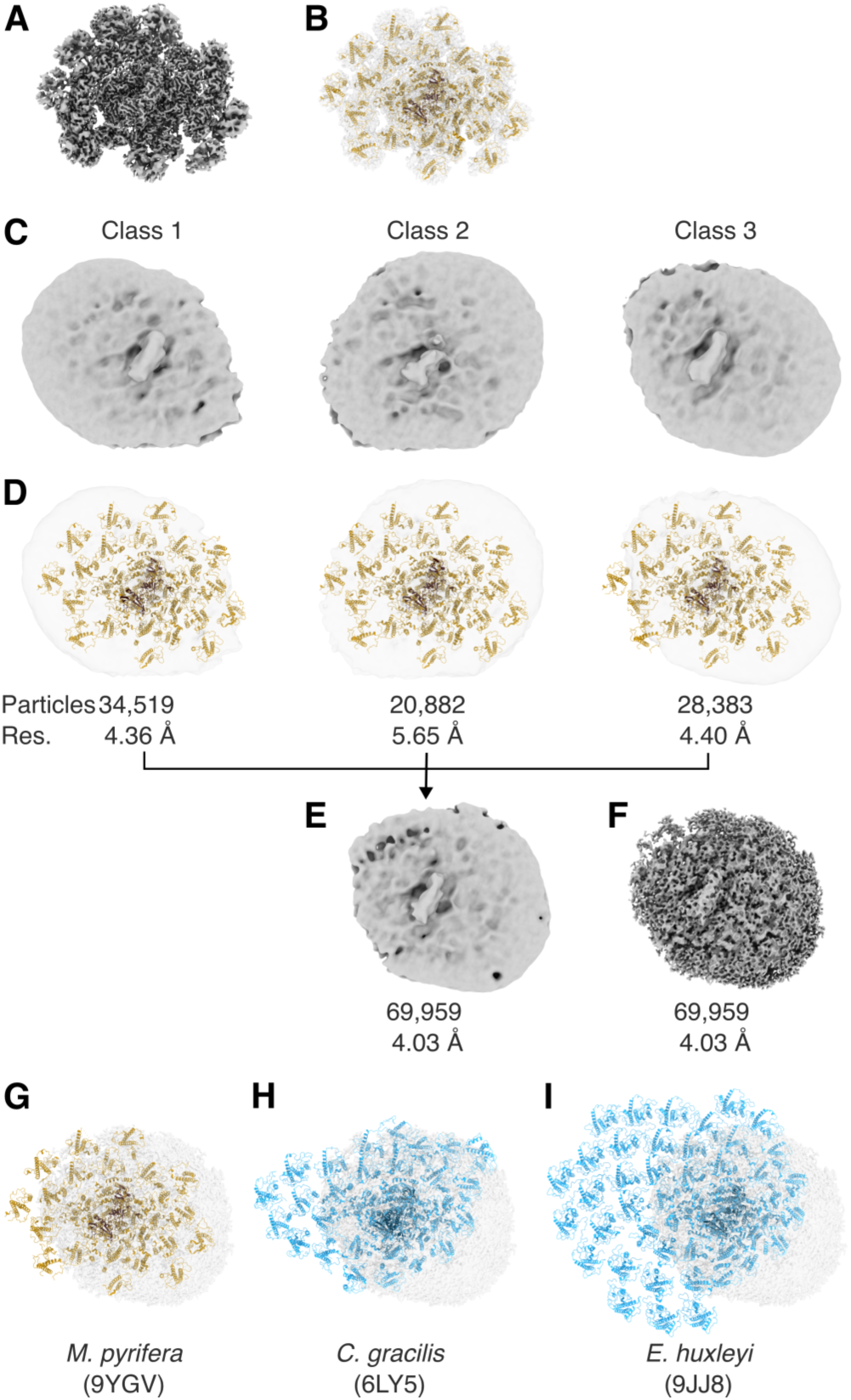
Larger PSI-FCP antenna from *M. pyrifera*. (**A-B**) Map (A) and model superposed on semitransparent map (B) of *M. pyrifera* PSI-FCP supercomplex discussed in all other figures or this work (PDB: 9YGV). Model shown in cartoon, colored in brown. Stromal domain (PsaC/D/E) colored in darker brown as visual anchor. (**C-F**) Larger PSI-FCP supercomplex obtained from *M. pyrifera* preparation extracted in different conditions relative to (A-B). (C-D) Reconstructions of three PSI-FCP supercomplex classes (class 1-3)obtained after non-uniform refinement. To reduce high-resolution noise and make the antenna architecture more evident, maps were Gaussian-filtered with 3 standard deviations. Maps shown alone (C) or with the *M. pyrifera* model (PDB: 9YGV, this work) fitted and superposed. Colored as in (B). Particle numbers and resolutions (Res.) for each class are indicated. (**E-F**) Reconstruction of the combined particles from classes 1-3 in (C-D) obtained after non-uniform refinement. Reconstruction shown with (E) or without (F) Gaussian filter as in (C-D). (**G-I**) Reconstruction from (F) fitted with superposed models of PSI-FCP supercomplexes from *M. pyrifera* (PDB: 9YGV, this work) (G), diatom *C. gracilis* (PDB: 6LY5) (*28*) (H) or *E. huxleyi* (PDB: 9JJ8) (*25*) (I). Models shown in cartoon and colored in brown with stromal domain in darker brown (G) or in blue with stromal domain in darker blue (H, I).

**Fig. S13.**
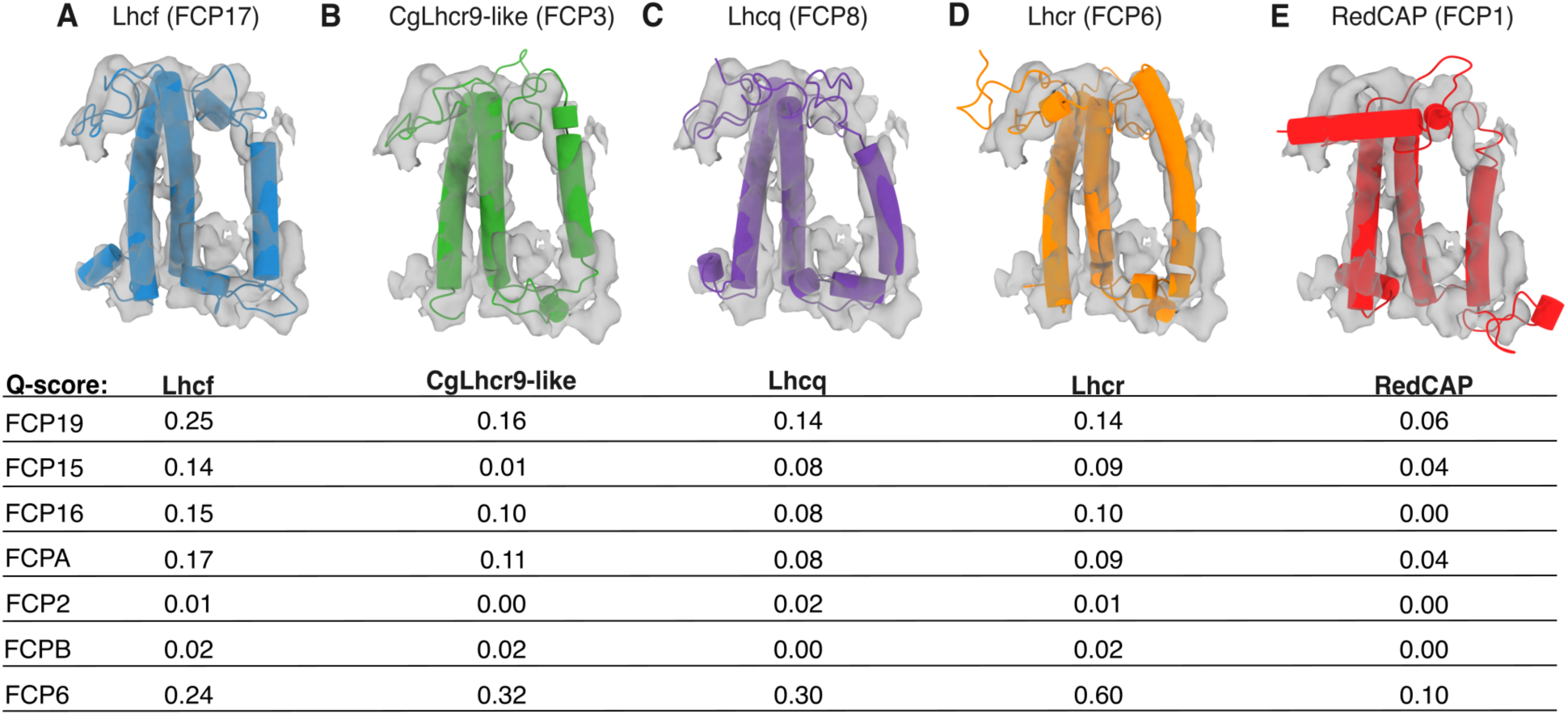
FCP subfamily assignment by map-model fit (Q score). (**A-E**) Representative example of *M. pyrifera* focused-refined map (FCPA shown) fit with representative FCPs for each assigned subfamily, arranged in decreasing Q-score fit. The model of *M. pyrifera* FCP6 was used for Lhcr, FCP3 for CgLhcr9-like, FCP1 for RedCAP, FCP8 for Lhcq, FCP17 for Lhcf. Tabulation below (A-E) shows full Q-scores for all un-assigned FPCs fit with all FCP subfamilies.

**Fig. S14.**
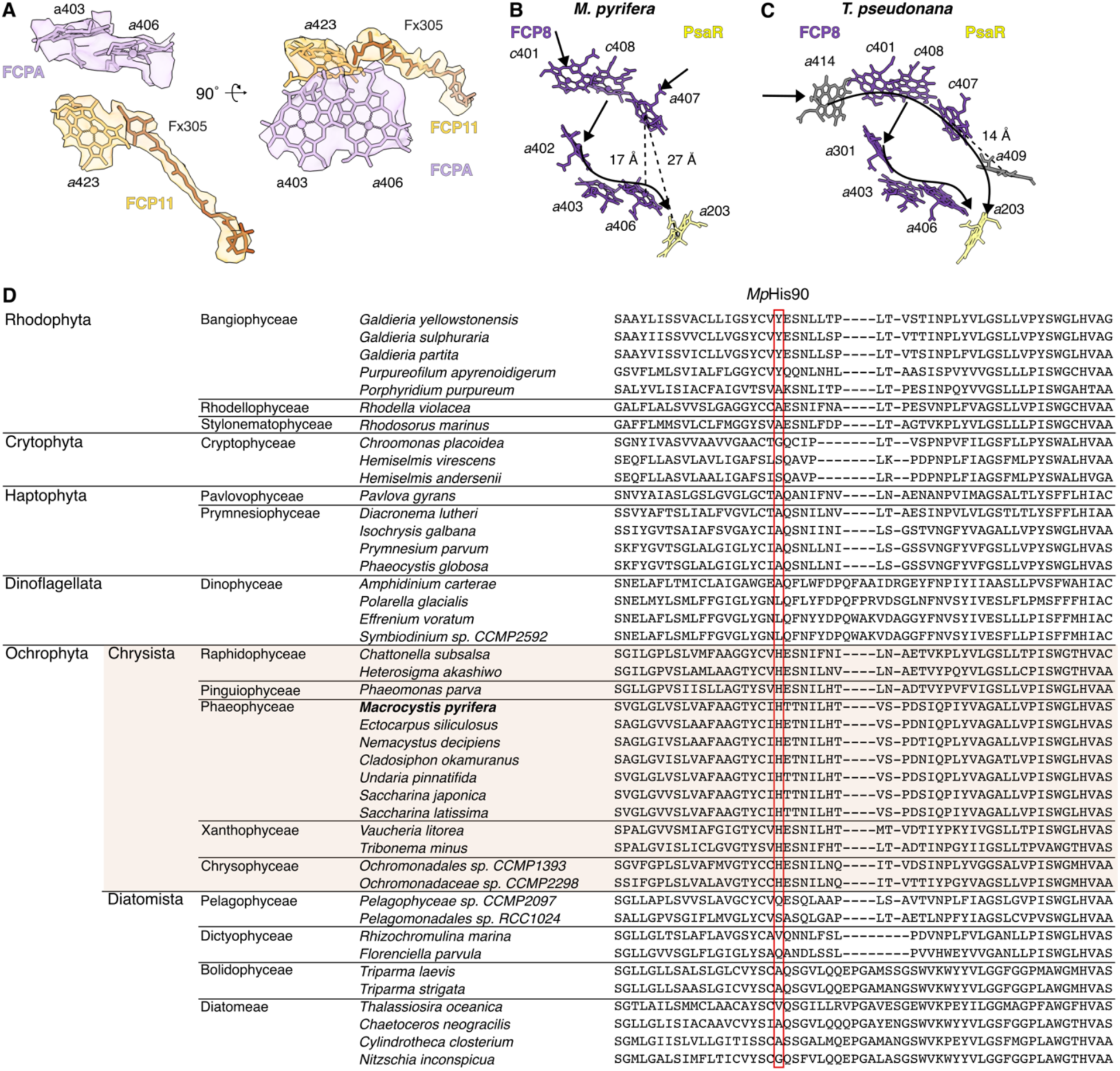
Additional details on chromophore features of *M. pyrifera*’s PSI-FCP. (**A**) Map-model fit of chlorophyll *a* molecules modeled in *M. pyrifera* FCPA and its presumed transfer partner in FCP11. Fucoxanthin (Fx) molecule discussed in text also shown. Models colored by subunit, map shown in semi-transparent surface colored by subunit. (**B**-**C**) Stromal connections in the FCP8/9-PsaR pathways of *M. pyrifera* (PDB: 9YGV, this work) (B) and diatom *T. pseudonana* (PDB: 8ZEH) (*31*) (C), viewed from the stroma. Chlorophyll molecules colored by subunit. Diatom FCP8 chlorophyll molecules missing in *M. pyrifera* colored in grey. Arrows denote predicted EET transfers. (**D**) Alignment of PsaR sequences from various ochrophytes with available genomes retrieved from NCBI GenBank, PhycoCosm and EukProt v3 databases, arranged by clade on the left. Homologous position for His90, which coordinates PsaR-chl *a*205 in *M. pyrifera*, marked with red border. Chrysista clade marked in light brown box.

**Fig. S15.**
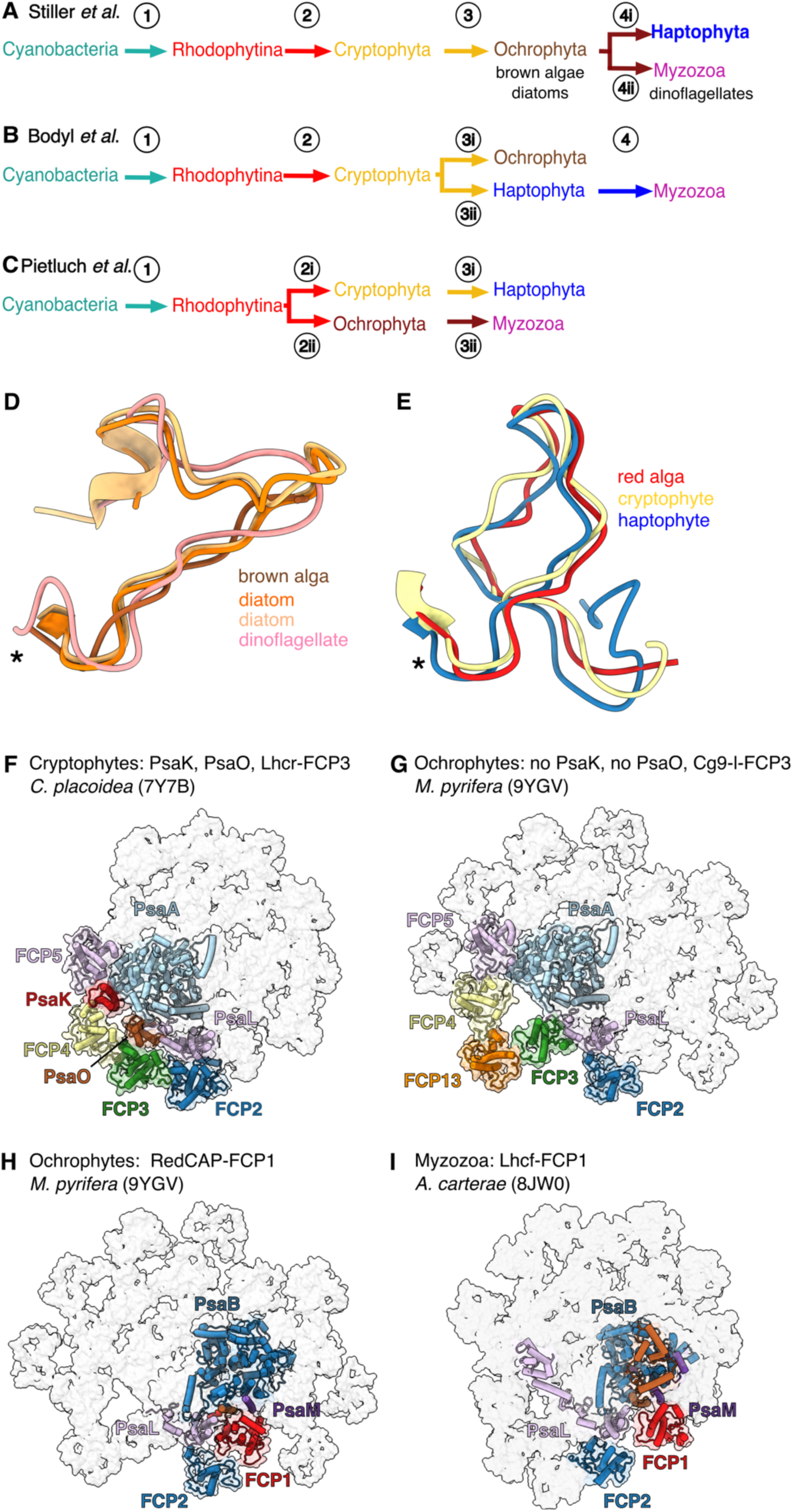
Models of serial endosymbiosis in red-derived organisms. (**A-C**) Models of endosymbioses in the red lineage consistent with molecular timescale analysis (*14*, *15*) proposed by Stiller *et al*. (*85*) (A), Bodyl *et al*. (*86*) (B) and Pietluch *et al.* (*15*) (C). Numbering in circles refers to levels of endosymbiosis. See **fig. S1D** for further details. (**D**-**E**) Superposition of FCP9 N-terminal region of red algae (*P. purpureum*, PDB: 7Y5E) (*19*), cryptophytes (*C. placoidea*, PDB: 7Y7B) (*22*) and haptophytes (*I. galbana*, PDB: 8Z11) (*24*) (F) or brown algae (*M. pyrifera*, PDB: 9YGV, this work), diatoms (*C. gracilis*, PDB: 6L4U) (*29*) and *T. pseudonana* (PDB: 8ZEH) (*31*), as well as dinoflagellates (*A. carterae*, PDB: 8JW0) (*26*). Note that FCP structures in (D-E) fully align in the region marked by an asterisk. (**F**-**G**) Overview of differences in the PsaK/L/O, FCP2/3/4 region between cryptophyte *C. placoidea* (F) and ochrophyte *M. pyrifera* (G), viewed from the stroma. Relevant subunits shown in colored cartoon over semi-transparent surface of the whole supercomplex. (**H-I**) Overview of differences in the FCP1 region, viewed from the stroma between ochrophyte *M. pyrifera* (H) and dinoflagellate (myzozoa) *A. carterae* (I). Relevant subunits shown in colored cartoon over semi-transparent surface of the whole supercomplex. PDB codes and references as in (D-E).

**Table S1.** CryoEM data collection and atomic model statistics.

| **Data collection and image processing** |  |
| --- | --- |
| Microscope | Titan Krios |
| Voltage (kV) | 300 |
| Camera | Falcon 4i |
| Data collection software | EPU |
| Electron exposure (e<sup>-</sup>/Å<sup>2</sup>) | 50 |
| Exposure time (s) | 4.95 |
| Frame number | 40 |
| Defocus range (μm) | -1 to -2 |
| Pixel size (Å) | 1.217 |
| Number of micrographs | 15690 |
| EMPIAR accession code | 12998 |
| Number of particles for final reconstruction | 133,000 |
| Symmetry imposed | C1 |
| Map Resolution range (Å) | 2.47-5.21 |
| FSC threshold | 0.143 |
| **Model refinement statistics** |  |
| Cross-correlation |  |
| Mask | 0.84 |
| Volume | 0.81 |
| Model composition |  |
| Non-hydrogen atoms | 53507 |
| Protein residues | 5047 |
| Ligands | 336 |
| Ramachandran |  |
| Favored (%) | 94.02 |
| Allowed (%) | 5.95 |
| Outlier (%) | 0.03 |
| Rotamer outliers (%) | 1.71 |
| Clash core | 6.82 |
| RMSD |  |
| Bond length (Å) | 0.007 |
| Bon angle (°) | 1.581 |
| B factors (Å<sup>2</sup>, min/max/mean) |  |
| Protein | 8.19/204.95/80.16 |
| Ligands | 12.08/218.80/78.48 |
| MolProbity score | 1.95 |
| PDB Accession code | 9YGV |

**Table S2.** Summary of deposited maps. Details of composite and focused refined maps deposited in EMDB database. FCP, fucoxanthin-chlorophyll-*a*/*c*-proteinMp, *Macrocystis pyrifera*; n.a., not applicable.

| Map | EMDB code | Particles | Res. (Å) | Map entry title |
| --- | --- | --- | --- | --- |
| Composite | 72770 | n.a. | 2.47-5.21 | Composite map of MpPSI-FCP supercomplex |
| Focused refined maps |  |  |  |  |
| PSI | 72805 | 133,000 | 2.47 | Focused refinement of MpPSI-FCP around PSI |
| FCP1 | 72806 | 133,000 | 2.95 | Focused refinement of MpPSI-FCP around FCP1 |
| FCP2 | 72807 | 22,780 | 4.71 | Focused refinement of MpPSI-FCP around FCP2 |
| FCP3 | 72808 | 133,000 | 3.1 | Focused refinement of MpPSI-FCP around FCP3 |
| FCP4 | 72809 | 104,446 | 3.7 | Focused refinement of MpPSI-FCP around FCP4 |
| FCP5 | 72810 | 121,591 | 3.17 | Focused refinement of MpPSI-FCP around FCP5 |
| FCP6 | 72811 | 133,000 | 3.18 | Focused refinement of MpPSI-FCP around FCP6 |
| FCP7 | 72812 | 133,000 | 3.11 | Focused refinement of MpPSI-FCP around FCP7 |
| FCP8 | 72813 | 133,000 | 2.97 | Focused refinement of MpPSI-FCP around FCP8 |
| FCP9 | 72814 | 93,065 | 3.78 | Focused refinement of MpPSI-FCP around FCP9 |
| FCP10 | 72816 | 133,000 | 3.14 | Focused refinement of MpPSI-FCP around FCP10 |
| FCP11 | 72817 | 119,214 | 3.02 | Focused refinement of MpPSI-FCP around FCP11 |
| FCP13 | 72818 | 104,924 | 3.82 | Focused refinement of MpPSI-FCP around FCP13 |
| FCP15 | 72819 | 74,423 | 4.71 | Focused refinement of MpPSI-FCP around FCP15 |
| FCP16 | 72820 | 94,294 | 4.81 | Focused refinement of MpPSI-FCP around FCP16 |
| FCP17 | 72821 | 92,002 | 3.66 | Focused refinement of MpPSI-FCP around FCP17 |
| FCP19 | 72822 | 28,903 | 4.92 | Focused refinement of MpPSI-FCP around FCP19 |
| FCPA | 72823 | 91,906 | 4.73 | Focused refinement of MpPSI-FCP around FCPA |
| FCPB | 72824 | 45,411 | 5.21 | Focused refinement of MpPSI-FCP around FCPB |

**Table S3.** Model statistics by subunit. *M. pyrifera* PSI-FCP supercomplex model details by subunit. PQN, phylloquinone; SF4, [4Fe-4S] cluster; CLA, chlorophyll *a*; KC1, CLA, chlorophyll *c*; A86, fucoxanthin; XAT, violaxanthin; BCR, beta-carotene; DGD, digalactosyl-diacyl-glycerol; LHG, di-palmitoyl-phosphatydil-glycerole; LMG, distearoyl-monogalactosyl-diglyceride; n.a., not available.

| Subunit name | Gene identifier | Gene database | Location | Chain ID | Total residues | Modelled residues |
| --- | --- | --- | --- | --- | --- | --- |
| **PsaA** | A0A8F0JYL2 | UniprotKB | PSI | A | 749 | 8-749 |
| **PsaB** | A0A8F0F8K8 | UniProtKB | PSI | B | 734 | 4-733 |
| **PsaC** | A0A8F0F8D0 | UniProtKB | PSI | C | 81 | 2-81 |
| **PsaD** | A0A8F0F8I9 | UniProtKB | PSI | D | 132 | 7-131 |
| **PsaE** | A0A8F0F8D1 | UniProtKB | PSI | E | 61 | 2-61 |
| **PsaF** | A0A8F0JY91 | UniProtKB | PSI | F | 203 | 43-203 |
| **PsaI** | A0A8F0F8C2 | UniProtKB | PSI | I | 36 | 2-35 |
| **PsaJ** | A0A8F0F8B4 | UniProtKB | PSI | J | 42 | 1-41 |
| **PsaL** | A0A8F0F8N6 | UniProtKB | PSI | L | 145 | 2-143 |
| **PsaM** | A0A8F0JYJ8 | UniProtKB | PSI | M | 30 | 1-30 |
| **PsaR** | CAM9152197.1 | GenBank | PSI | R | 132 | 48-132 |
| **FCP1** | 9470359 | PhycoCosm | 1<sup>st</sup> belt | 1 | 231 | 54-231 |
| **FCP2** | unassigned | n.a. | 1<sup>st</sup> belt | 2 | n.a. | n.a. |
| **FCP3** | 9638407 | PhycoCosm | 1<sup>st</sup> belt | 3 | 191 | 27-181 |
| **FCP4** | DN157149 | PhycoCosm | 1<sup>st</sup> belt | 4 | 237 | 40-235 |
| **FCP5** | 9353087 | PhycoCosm | 1<sup>st</sup> belt | 5 | 222 | 44-218 |
| **FCP6** | 9608268 | PhycoCosm | 1<sup>st</sup> belt | 6 | 212 | 39-211 |
| **FCP7** | DN156408 | PhycoCosm | 1<sup>st</sup> belt | 7 | 169 | 1-169 |
| **FCP8** | 9480344 | PhycoCosm | 1<sup>st</sup> belt | 8 | 215 | 40-214 |
| **FCP9** | 9541138 | PhycoCosm | 1<sup>st</sup> belt | 9 | 216 | 62-208 |
| **FCP10** | 9608268 | PhycoCosm | 1<sup>st</sup> belt | 10 | 212 | 40-211 |
| **FCP11** | 9689568 | PhycoCosm | 1<sup>st</sup> belt | 11 | 213 | 45-211 |
| **FCP13** | 9541138 | PhycoCosm | 2<sup>nd</sup> belt | 13 | 216 | 66-170,<br>176-214 |
| **FCP15** | unassigned | n.a. | 2<sup>nd</sup> belt | 15 | n.a. | n.a. |
| **FCP16** | unassigned | n.a. | 2<sup>nd</sup> belt | 16 | n.a. | n.a. |
| **FCP17** | 1228943 | PhycoCosm | 2<sup>nd</sup> belt | 17 | 218 | 46-149,<br>164-205 |
| **FCP19** | unassigned | n.a. | 2<sup>nd</sup> belt | 19 | n.a. | n.a. |
| **FCPA** | unassigned | n.a. | 2<sup>nd</sup> belt | a | n.a. | n.a. |
| **FCPB** | unassigned | n.a. | 2<sup>nd</sup> belt | b | n.a. | n.a. |

**Table S3 (continued). Model statistics by subunit.**
| Subunit name | PQN | SF4 | CLA | KC1 | A86 | XAT | BCR | DGD | LHG | LMG |
| --- | --- | --- | --- | --- | --- | --- | --- | --- | --- | --- |
| **PSI** |  |  |  |  |  |  |  |  |  |  |
| PsaA | 1 | 1 | 44 | - | 1 | - | 5 | - | 5 | - |
| PsaB | 1 | - | 40 | - | 1 | - | 6 | - | 2 | - |
| PsaC | - | 2 | - | - | - | - | - | - | - | - |
| PsaD | - | - | - | - | - | - | - | - | - | - |
| PsaE | - | - | - | - | - | - | - | - | - | - |
| PsaF | - | - | 4 | - | - | - | 2 | - | - | - |
| PsaI | - | - | 1 | - | - | - | 2 | - | - | - |
| PsaJ | - | - | 1 | - | - | 1 | 1 | - | - | 1 |
| PsaL | - | - | 3 | - | - | - | 2 | - | - | - |
| PsaM | - | - | - | - | - | - | 1 | - | - | - |
| PsaR | - | - | 2 | - | 1 | - | 1 | 1 | - | - |
| Total PSI | 2 | 3 | 95 | 0 | 3 | 1 | 20 | 1 | 7 | 1 |
| **FCP** |  |  |  |  |  |  |  |  |  |  |
| FCP1 | - | - | 7 | 1 | 2 | 1 | 1 | - | 1 | - |
| FCP2 | - | - | 5 | - | - | - | - | - | - | - |
| FCP3 | - | - | 6 | 3 | 3 | 1 | - | - | - | - |
| FCP4 | - | - | 7 | 4 | 2 | 2 | - | 1 | - | - |
| FCP5 | - | - | 8 | 1 | 2 | 2 | - | 1 | - | - |
| FCP6 | - | - | 8 | 4 | 1 | 4 | - | - | 1 | - |
| FCP7 | - | - | 9 | 2 | - | 5 | - | - | 1 | - |
| FCP8 | - | - | 7 | 2 | 1 | 2 | 1 | - | - | - |
| FCP9 | - | - | 7 | 2 | 1 | 1 | - | - | 1 | - |
| FCP10 | - | - | 9 | 3 | 2 | 3 | - | - | 2 | - |
| FCP11 | - | - | 9 | 1 | 1 | 1 | - | - | 1 | - |
| FCP13 | - | - | 6 | 1 | - | 2 | - | - | - | - |
| FCP15 | - | - | 4 | - | - | - | - | - | - | - |
| FCP16 | - | - | 6 | - | - | - | - | - | - | - |
| FCP17 | - | - | 7 | 2 | 1 | 1 | 1 | - | - | - |
| FCP19 | - | - | 5 | - | - | - | - | - | - | - |
| FCPA | - | - | 8 | 1 | - | - | - | - | - | - |
| FCPB | - | - | 5 | - | - | - | - | - | - | - |
| Total FCP | 0 | 0 | 123 | 27 | 16 | 25 | 3 | 2 | 7 | 0 |
| Total PSI-FCP | 2 | 3 | 218 | 27 | 19 | 26 | 23 | 3 | 14 | 1 |

**Table S4.** Correspondence between chromophore PDB numbering and binding site numbering. Binding site numbering per (*31*)*. a*, chlorophyll *a*; *c*, chlorophyll *c*; Bcr, beta-carotene; Fx, fucoxanthin; Vx, violaxanthin.

| Site | FCP1 | FCP2 | FCP3 | FCP4 | FCP5 | FCP6 | FCP7 | FCP8 | FCP9 | FCP10 |
| --- | --- | --- | --- | --- | --- | --- | --- | --- | --- | --- |
| 301 | - | - | Fx313 | - | - | Vx321 | Vx322 | - | - | Fx326 |
| 302 | Fx301 | - | Fx314 | Vx340 | Vx321 | Vx325 | Vx323 | Vx319 | Fx315 | Vx321 |
| 303 | Fx318 | - | Vx<br>317 | Vx<br>343 | Fx312 | Vx324 | Vx321 | Vx318 | Vx317 | Vx323 |
| 304 | - | - | - | - | - | - | - | - | - | - |
| 305 | Bcr317 | - | Fx311 | Fx335 | Fx322 | Fx323 | Vx320 | Fx311 | - | Vx322 |
| 306 | - | - | - | Fx338 | - | Vx320 | Vx324 | - | - | Fx319 |
| 307 | - | - | - | - | Vx320 | - | - | Bcr315 | - | - |
| 308 | Vx316 | - | - | - | - | - | - | - | - | - |
| 401 | - | - | *a*307 | *c*326 | *c*316 | *a*307 | *a*308 | *c*306 | - | *a*309 |
| 402 | *a*305 | - | *a*301 | *a*321 | *a*302 | *a*301 | *a*301 | *a*301 | *a*301 | *a*303 |
| 403 | *a*312 | *a*301 | *c*302 | *a*322 | *a*303 | *a*302 | *a*302 | *a*302 | *a*302 | *a*304 |
| 404 | - | - | *a*303 | *a*323 | *a*304 | *a*303 | *a*303 | *a*303 | *a*303 | *a*305 |
| 405 | *a*313 | *a*303 | *a*304 | - | *a*306 | *a*304 | *a*318 | - | *c*304 | *a*306 |
| 406 | *a*308 | *a*304 | *a*305 | *a*324 | *a*305 | *c*305 | *a*306 | *a*304 | *a*305 | *a*307 |
| 407 | *a*309 | *a*305 | *a*306 | *a*325 | *a*324 | *a*306 | *a*307 | *a*305 | *a*306 | *a*308 |
| 408 | *c*310 | - | *c*308 | *c*327 | *a*318 | *c*308 | *c*309 | *a*307 | *c*308 | *a*310 |
| 409 | *a*300 | - | *c*309 | *a*328 | - | *c*309 | *a*310 | *a*320 | *a*309 | *a*311 |
| 410 | *a*315 | - | - | *c*331 | - | *a*311 | - | - | *a*310 | *a*325 |
| 412 | - | - | - | - | - | *a*322 | *a*319 | - | - | *a*314 |
| 413 | - | - | - | - | - | - | - | *a*316 | - | - |
| 414 | - | - | - | *c*330 | - | - | - | - | - | - |
| 415 | - | - | - | *a*341 | *a*310 | *c*310 | *c*311 | - | - | *c*312 |
| 416 | - | - | - | - | - | - | - | - | - | - |
| 417 | - | - | - | - | - | - | - | - | - | - |
| 419 | - | - | - | - | - | - | - | - | - | - |
| 420 | - | - | - | - | - | - | - | - | - | - |
| 421 | - | - | - | - | - | - | - | - | - | - |
| 422 | - | - | - | - | - | - | - | - | - | - |
| 423 | - | - | - | - | - | - | - | - | - | - |

**Table S4 (continued). Correspondence between chromophore PDB numbering and binding site numbering.** Binding site numbering per (31). *a*, chlorophyll *a*; *c*, chlorophyll *c*; Bcr, beta-carotene; A86, fucoxanthin; Vx, violaxanthin.
| Site | FCP11 | FCP13 | FCP17 | FCP15 | FCP16 | FCP19 | FCPA | FCPB |
| --- | --- | --- | --- | --- | --- | --- | --- | --- |
| 301 | - | - | - | - | - | - | - | - |
| 302 | - | Vx317 | Fx315 | - | - | - | - | - |
| 303 | Vx316 | Vx316 | Vx318 | - | - | - | - | - |
| 304 | - | - | - | - | - | - | - | - |
| 305 | Fx309 | - | Bcr317 | - | - | - | - | - |
| 306 | - | - | - | - | - | - | - | - |
| 307 | - | - | - | - | - | - | - | - |
| 308 | - | - | - | - | - | - | - | - |
| 401 | *a*305 | - | *a*309 | - | - | - | *a*299 | - |
| 402 | *a*301 | *a*301 | *c*319 | *a*269 | *a*307 | *a*261 | *a*300 | *a*228 |
| 403 | *a*302 | *a*302 | *a*303 | - | *a*308 | *a*262 | *a*301 | *a*229 |
| 404 | *a*312 | *a*303 | *a*304 | *a*271 | - | - | *a*302 | *a*230 |
| 405 | - | *c*304 | *a*316 | *a*272 | - | - | *a*303 | - |
| 406 | *a*303 | *a*305 | *a*307 | - | - | *a*266 | *a*304 | - |
| 407 | *a*304 | *a*306 | *a*308 | *a*274 | - | *a*267 | *a*305 | - |
| 408 | *a*315 | - | *c*310 | - | *a*315 | - | *c*306 | - |
| 409 | *a*307 | - | *a*311 | - | *a*314 | *a*270 | *a*307 | *a*236 |
| 410 | - | *a*310 | - | - | - | - | - | - |
| 412 | - | - | - | - | - | - | - | - |
| 413 | - | - | - | - | - | - | - | - |
| 414 | - | - | - | - | - | - | - | - |
| 415 | *c*308 | - | - | - | - | - | - | - |
| 416 | - | - | - | - | - | - | - | - |
| 417 | - | - | - | - | - | - | - | - |
| 419 | - | - | - | - | - | - | - | - |
| 420 | - | - | - | - | - | - | - | - |
| 421 | - | - | - | - | - | - | - | - |
| 422 | - | - | - | - | - | - | - | - |
| 423 | *a*311 | - | - | - | - | - | - | - |

**Table S5.**
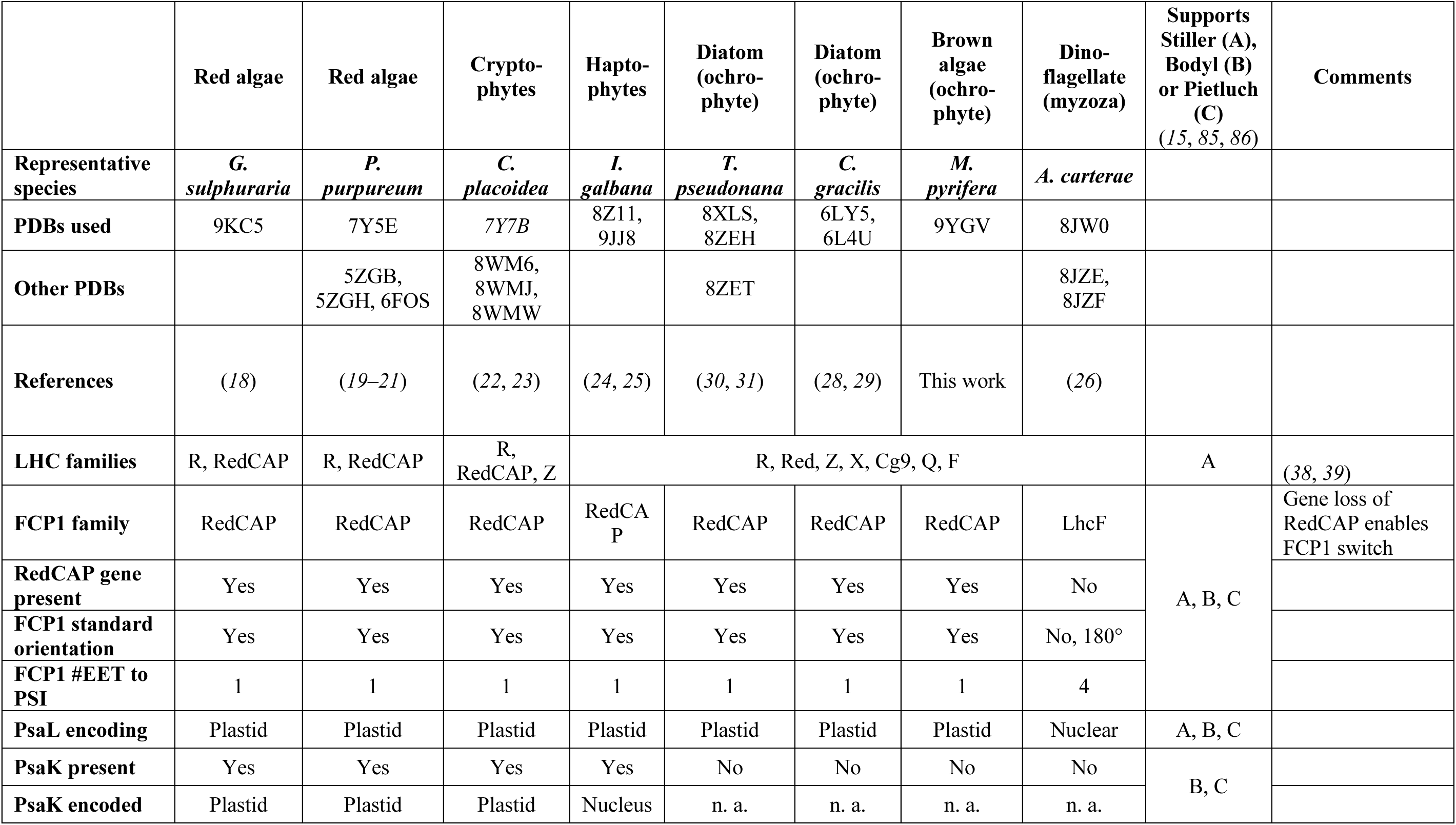
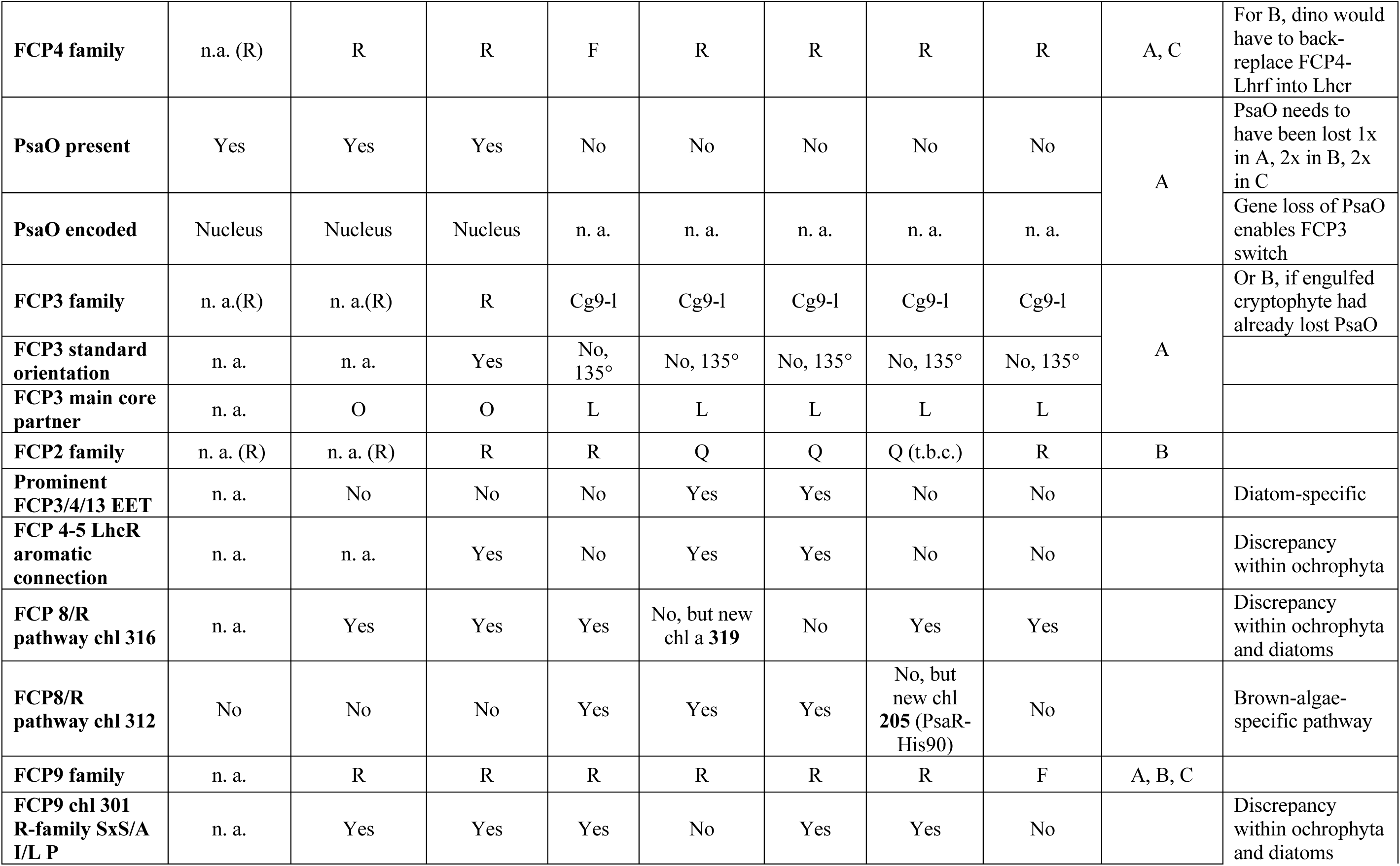
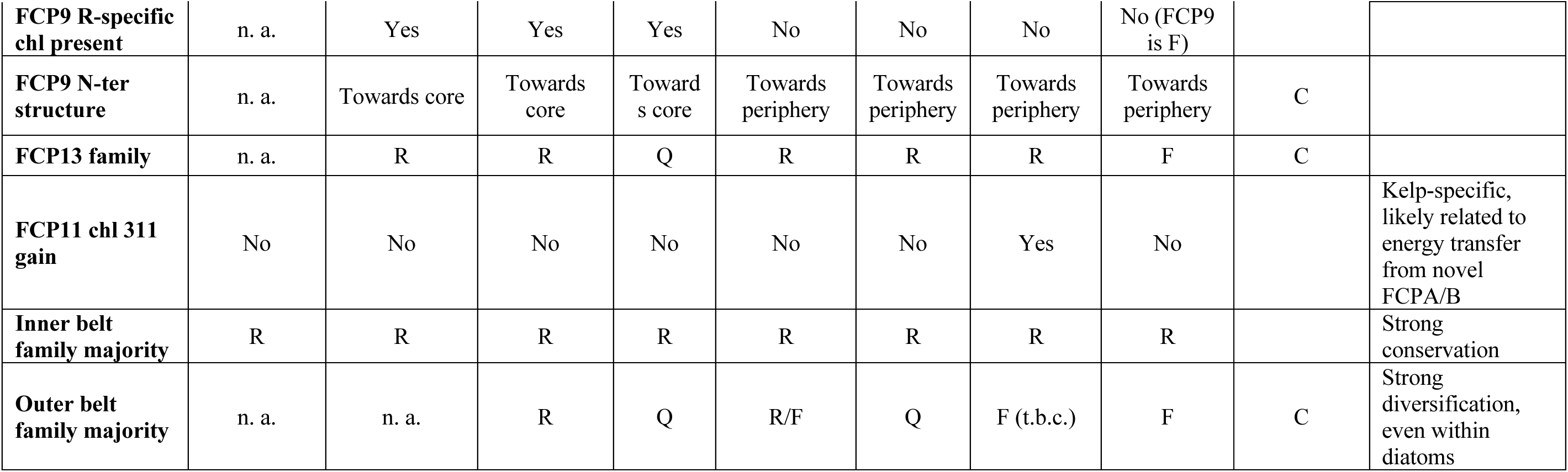
Summary of differences between red-lineage clades. Summary of clade differences in protein and chromophore features discussed in text. FCP subfamilies represented by the last letter, e.g., R, Lhcr. Cg9-l, CgLhcr9-like; n.a., not applicable; t.b.c., to be confirmed.

Submitted Manuscript: Confidential

Template revised July 2024

**Data File 1. Sequences of FCP and related LHC proteins used in phylogenetic analysis.** FCP sequences for *Macrocystis pyrifera*, *Ectocarpus siliculosus*, *Chaetoceros gracilis*, *Thalassiosira pseudonana* and green algal LHC proteins used to root the trees in **fig. S9**.

**Data File 2. Membrane thickness of FCP and core helices across the red lineage. Data File 2. Membrane thickness of FCP and core helices across the red lineage**. Measurements of the membrane span of the PsaA/B and FCP helices A-C for subfamilies Lhcr, Lhcq, Lhcq-like, Lhcf measured from *Macrocystis pyrifera* (PDB: 9YGV, this work), *Emiliania huxleyi* (PDB: 9JJ8) (*25*) and *Galdieria sulphuraria* (PDB: (9KC5) (*18*) shown in **Fig. 2**. To determine the span of the helices, measurements were taken using the distance tool in Coot and values were scaled relative to the field of view of the membrane in Inkscape. To determine the averages for each FCP subfamily, at least 3 measurements were taken for each helix of the different FCP subfamilies; when possible, at least three subfamily members were measured.

